# Evolution and insights into the structure and function of the DedA superfamily containing TMEM41B and VMP1

**DOI:** 10.1101/2020.12.18.423352

**Authors:** Fumiya Okawa, Yutaro Hama, Sidi Zhang, Hideaki Morishita, Hayashi Yamamoto, Tim P. Levine, Noboru Mizushima

**Affiliations:** Department of Biochemistry and Molecular Biology, Graduate School of Medicine, The University of Tokyo, Tokyo 113-0033, Japan; UCL Institute of Ophthalmology, University College London, London EC1V 9EL, UK; Department of Physiology, Juntendo University Graduate School of Medicine, Tokyo 113-8421, Japan

**Keywords:** DedA, VMP1, TMEM41B, reentrant loop, autophagy

## Abstract

TMEM41B and VMP1 are endoplasmic reticulum (ER)-localizing multi-spanning membrane proteins required for ER-related cellular processes such as autophagosome formation, lipid droplet homeostasis, and lipoprotein secretion in eukaryotes. Both proteins have a VTT domain, which is similar to the DedA domain found in bacterial DedA family proteins. However, the molecular function and structure of the DedA and VTT domains (collectively referred to as DedA domains) and the evolutionary relationships among the DedA domain-containing proteins are largely unknown. Here, we conduct remote homology search and identify a new clade consisting mainly of bacterial PF06695 proteins of unknown function. Phylogenetic analysis reveals that the TMEM41, VMP1, DedA, and PF06695 families form a superfamily with a common origin, which we term the DedA superfamily. Coevolution-based structural prediction suggests that the DedA domain contains two reentrant loops that face each other in the membrane. This topology is biochemically verified by the substituted cysteine accessibility method. The predicted structure is topologically similar to that of the substrate-binding region of Na^+^-coupled glutamate transporter solute carrier 1. A potential ion-coupled transport function of the DedA superfamily proteins is discussed.

## INTRODUCTION

TMEM41B and VMP1 are endoplasmic reticulum (ER)-localizing multi-spanning membrane proteins essential for autophagosome formation, lipid droplet homeostasis, membrane contact, and lipoprotein secretion in metazoans (Demignot et al., 2014; Moretti et al., 2018; Morishita et al., 2019; Morita et al., 2018; Ropolo et al., 2007; Shoemaker et al., 2019; Tabara and Escalante, 2016; Zhao et al., 2017). As lipid droplets and lipoproteins are directly generated from the ER (Demignot et al., 2014; Walther et al., 2017) and autophagosomes are also formed on the ER (Nakatogawa, 2020), these proteins are considered to play fundamental roles in the ER. Elucidation of the molecular functions of TMEM41B and VMP1 would provide important insights into our understanding of the role of the ER in autophagy and other pathways, but their functions and structure are largely unknown.

VMP1 and TMEM41B contain a conserved transmembrane domain that is also found in TMEM64 and its homolog Tvp38 in metazoans, yeasts (Inadome et al., 2007), chloroplasts, and cyanobacteria (Keller and Schneider, 2013). We previously termed this domain the VTT (<u>V</u>MP1, <u>T</u>MEM41 and <u>T</u>vp38/TMEM64) domain (also known as SNARE_assoc domain; Pfam PF09335) (Morita et al., 2018; Morita et al., 2019). The VTT domain is similar to the bacterial <u>d</u>ownstream *<u>E</u>. coli* <u>D</u>NA gene <u>A</u> (from *hisT*) (DedA) domain (Doerrler et al., 2013; Inadome et al., 2007; Khafizov et al., 2010; Nonet et al., 1987; Thompkins et al., 2008). The DedA domain is present in a set of bacterial proteins that constitute the DedA family (Thompkins et al., 2008). YqjA and YghB are the best-characterized members of this family and are known to regulate temperature sensitivity, cell division (Thompkins et al., 2008), the export of periplasmic amidases (Sikdar and Doerrler, 2010), drug resistance, (Kumar and Doerrler, 2014), pH sensitivity (Kumar and Doerrler, 2015), and lipid composition of the cell membrane (Boughner and Doerrler, 2012; Thompkins et al., 2008). However, although putative transporter functions have been hypothesized based on genetic studies (Doerrler et al., 2013; Kumar and Doerrler, 2014), the molecular functions of these bacterial DedA family proteins are unknown. The VTT and DedA domains of most proteins in this family contain the conserved sequence motifs [F/Y]XXX[R/K] and GXXX[V/I/L/M]XXXX[F/Y] (Doerrler et al., 2013; Keller and Schneider, 2013; Tabara et al., 2019). While the VTT and DedA domains are evolutionarily related, previous phylogenetic analyses of the VTT and DedA domain-containing proteins were conducted with relatively small numbers of these proteins, excluding potential remote homologs (Boughner and Doerrler, 2012; Doerrler et al., 2013; Keller and Schneider, 2013; Thompkins et al., 2008). Thus, the exact definition of the VTT and DedA domains and their evolutionary relationships remain unclear.

Little is known about the structure of the VTT and DedA domains. The VTT domain was predicted to form a complicated structure containing several transmembrane helices (TMHs), some of which may be discontinuous (Morita et al., 2018; Morita et al., 2019). The DedA domain was proposed to adopt a structure similar to one half of the leucine transporter LeuT (Keller et al., 2014; Khafizov et al., 2010), leading to the hypothesis that the DedA domain serves as a half transporter module. Consistently, the self-interaction of YqjA has been reported (Keller et al., 2015). However, there is no experimental evidence supporting this structural prediction.

Here, we provide novel insights into the evolution and molecular functions of the VTT and DedA domains from both an expanded phylogenetic analysis that includes the remote homologs and a coevolution-based structural prediction. We found that the VTT and DedA domain-containing proteins, including the newly identified remote homolog Pfam PF06695, constitute a large superfamily with a common origin, which we term the DedA superfamily. The new phylogenetic tree suggests that prokaryotes already had several distinct ancestors, some of which evolved into the present eukaryotic homologs. Structure predictions and accompanying biochemical verifications define the membrane topology of the VTT/DedA domain, which contains two canonical TMHs and two reentrant loops that face each other in the membrane. Such structures are observed in transporters, ion channels, and a lipid dephosphorylating enzyme, suggesting a potential ion-coupled transporter-like function and a lipid-binding property for the DedA domain.

## RESULTS

### VTT/DedA domain-containing proteins form the DedA superfamily

To expand the phylogenetic analysis of the VTT and DedA domain-containing proteins, remote homology search was conducted using HHsearch, a hidden-Markov-model (HMM)-based method suitable for identifying homologous genes over long evolutionary distances (Steinegger et al., 2019). HHsearch considers both primary sequences and secondary structures, making it more sensitive when primary sequences have diverged among distantly related taxa, such as between eukaryotes and prokaryotes. Using human VMP1 as a query, we identified 125 homologous sequences in 23 species comprising eukaryotes, bacteria, and archaea (104 sequences after removing redundancy; Fig. S1). These sequences include all known proteins containing the VTT domain (TMEM41A, TMEM41B, TMEM64, Tvp38, YdjX, and YdjZ, the last two of which are also in the DedA family (Morita et al., 2018)) and all eight *E. coli* DedA family proteins (YdjX, YdjZ, YabI, DedA, YohD, YghB, YqjA, and YqaA) (Boughner and Doerrler, 2012; Doerrler et al., 2013), and this is consistent with a previous report that Tvp38 and the DedA family proteins are homologs (Keller and Schneider, 2013). The search also identified a new Pfam family, PF06695 putative small multi-drug export proteins. The majority of the members in this Pfam family are from bacteria, and their functions are unknown. A similar result (that PF06695 is a remote homolog) was obtained by using a PSI-blast-HHsearch combination (instead of HHblits-HHsearch), and by turning off the secondary structure scoring option in HHsearch (data not shown). Indeed, even PSI-blast alone revealed the remote homology (data not shown).

Alignment of representative sequences from the VTT and DedA domain-containing proteins including PF06695 (Fig. 1) shows that the homologous region extends beyond the previously suggested VTT domain (Morita et al., 2018) towards the N terminus by about 30 amino acids. The extended region is predicted to form a helix-coil-helix structure (“reentrant loop1” as described later). Of all the homologous sequences identified by HHsearch, we found that five bacterial proteins (YqaA, NP_388110 in *Bacillus subtilis*, YP_002348660 and YP_002348198 in *Yersinia pestis*, and NP_273579 in *Neisseria meningitidis*) and one archaeal protein (WP_048046344 in *Methanosarcina mazei*) consisted almost entirely of this aligned region alone, suggesting that this region could be a functional unit. Consistent with previous reports (Keller and Schneider, 2013; Tabara et al., 2019), the two motifs [F/Y]XXX[R/K] (motif 1) and GXXX[V/I/L/M]XXXX[F/Y] (motif 2) are conserved in VMP1, Tvp38, and most of the *E. coli* DedA family proteins, but not in TMEM41A/B, TMEM64, or the newly identified PF06695. For HsTMEM41A/B, motif 1 ends with tyrosine or serine and motif 2 starts with proline. For HsTMEM64, motif 1 starts with histidine and motif 2 starts with serine. For PF06695, motif 2 starts with asparagine and ends with alanine. Taken together, these results show that, despite minor differences, these proteins may form a large superfamily with a common origin, and we will hereafter refer to this superfamily as the DedA superfamily and the shared domain including the extended region as the DedA domain (an extended version of the previously defined VTT domain) (Fig. 1).

**Fig. 1.**
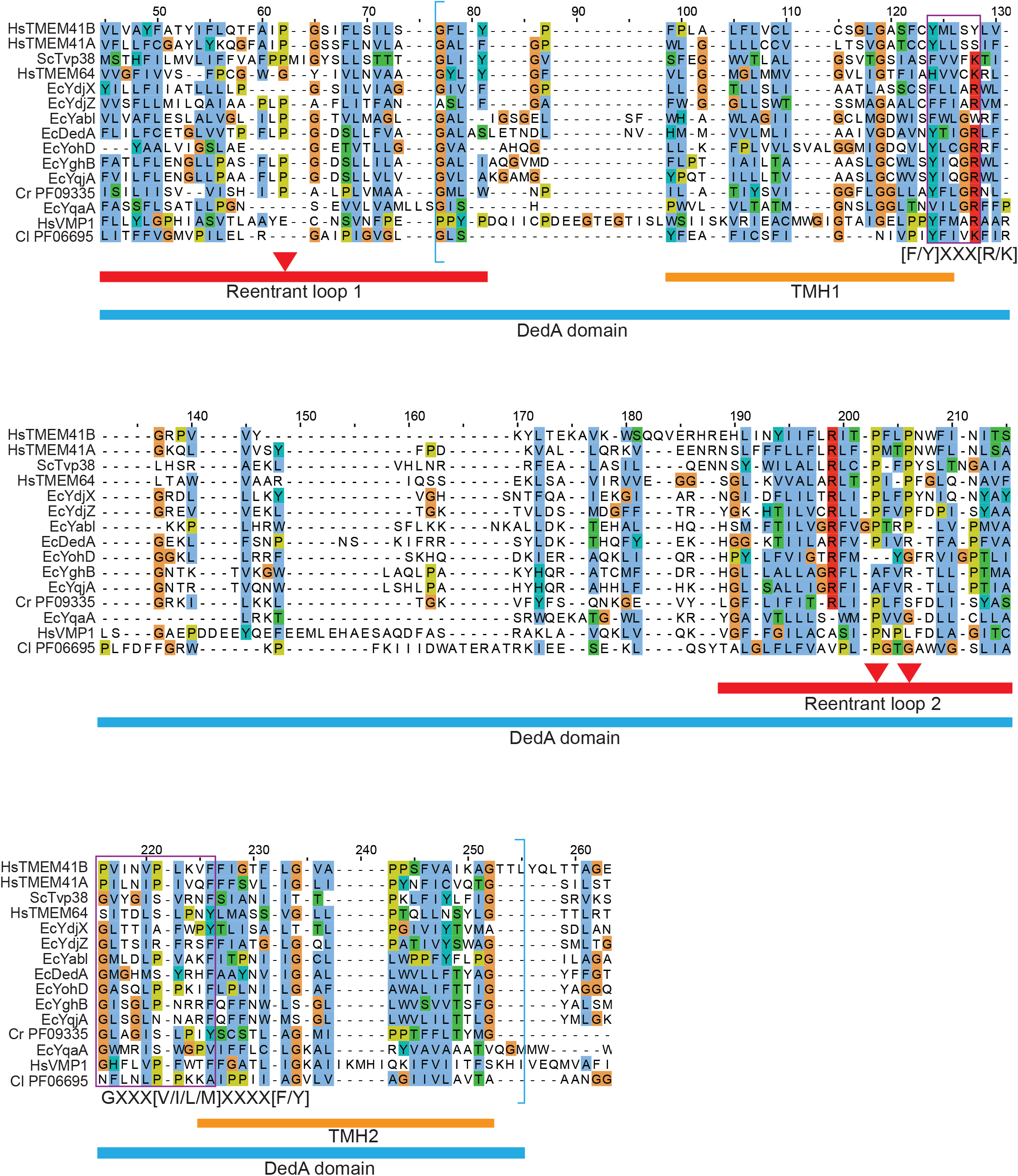
Multiple sequence alignment of representative sequences from the DedA superfamily. The numbers at the top are column numbers in the alignment. At the bottom, the blue bar shows the range of the DedA domain consisting of reentrant loop 1 and the previously proposed VTT domain (indicated by blue brackets around the sequences) (Morita et al., 2018), red bars show the ranges of reentrant loops with positions of conserved prolines flagged as triangles, and orange bars show the ranges of canonical transmembrane helices (TMHs) predicted by TMHMM. Purple boxes around the sequences show conserved motifs (Keller and Schneider, 2013; Tabara et al., 2019). Hs, *Homo sapiens*; Sc, *Saccharomyces cerevisiae*; Ec, *Escherichia coli*; Cr, *Crocosphaera subtropica*; Cl, *Clostridium species*. Cr PF09335 (SNARE_assoc) and Cl PF06695 (Small_multidrug) are representative sequences from PF09335 and PF06695, respectively.

To establish the evolutionary relationships among the DedA superfamily proteins, including the newly identified remote homologs, we reconstructed a phylogenetic tree using Graph Splitting (Matsui and Iwasaki, 2020) (Fig. 2A). This method outperforms classical methods such as maximum likelihood and Bayesian inference (Felsenstein, 1981; Rannala and Yang, 1996) when sequences are divergent, as it relies on all-to-all pairwise alignment instead of multiple sequence alignment, which shrinks significantly when sequence similarity is low. In the resulting phylogenetic tree, there are four families: the VMP1 family, a family including TMEM41A, TMEM41B, TMEM64, and Tvp38 (referred to as the TMEM41 family hereafter), the DedA family except for YdjX and YdjZ, and the PF06695 family. Note that Tvp38, which contains the two aforementioned sequence motifs, resides with TMEM41A, TMEM41B, and TMEM64, which are devoid of the two motifs, in the TMEM41 family. Bacterial YdjX and YdjZ are in the TMEM41 family, suggesting that they may be evolutionarily closer to the eukaryotic TMEM41 family proteins than to other DedA proteins, in agreement with an earlier report (Keller and Schneider, 2013). VMP1 is the outmost family in the eukaryotic cluster surrounded by the DedA and PF06695 families. Most eukaryotic proteins were found only in the TMEM41 and VMP1 families, but a few plant chloroplast proteins and SAR proteins were also found in the DedA (*Arabidopsis thaliana* NP_193051 and *Solanum lycopersicum* XP_004247084; #66 and #78 in Fig. 2B) and PF06695 (*Arabidopsis thaliana* NP_178363, *Solanum lycopersicum* XP_004238763 and *Thalassiosira oceanica* K0TKX5; #5, #73 and #111 in Fig, 2B) families. Because these proteins appear to exist in only limited lineages, they might have been transferred from the chloroplast to the nuclear genome after these lineages separated from other eukaryotes. Bacterial and archaeal proteins were found in all four families. Among homologous sequences from *Candidatus Prometheoarchaeum syntrophicum*, an archaeon very close to the branching point of archaea and eukaryotes (Imachi et al., 2020), one sequence (“seq2”) lies at the center of the TMEM41 family, another sequence (“seq1”) appears in the VMP1 family (Fig. 2A), and a third sequence (“seq3”) is at the periphery of the TMEM41 family, suggesting that there were probably different prokaryotic ancestors for the two eukaryotic families. These patterns can also be seen directly from the sequence similarity network behind the phylogenetic tree (Fig. 2B; Table S1), where similar sequences are clustered together. In summary, the phylogenetic analysis further supports the existence of the DedA superfamily and suggests that PF06695 probably branched out early, with ancestral DedA proteins splitting into three groups and developing into the DedA, VMP1, and TMEM41 families.

**Fig. 2.**
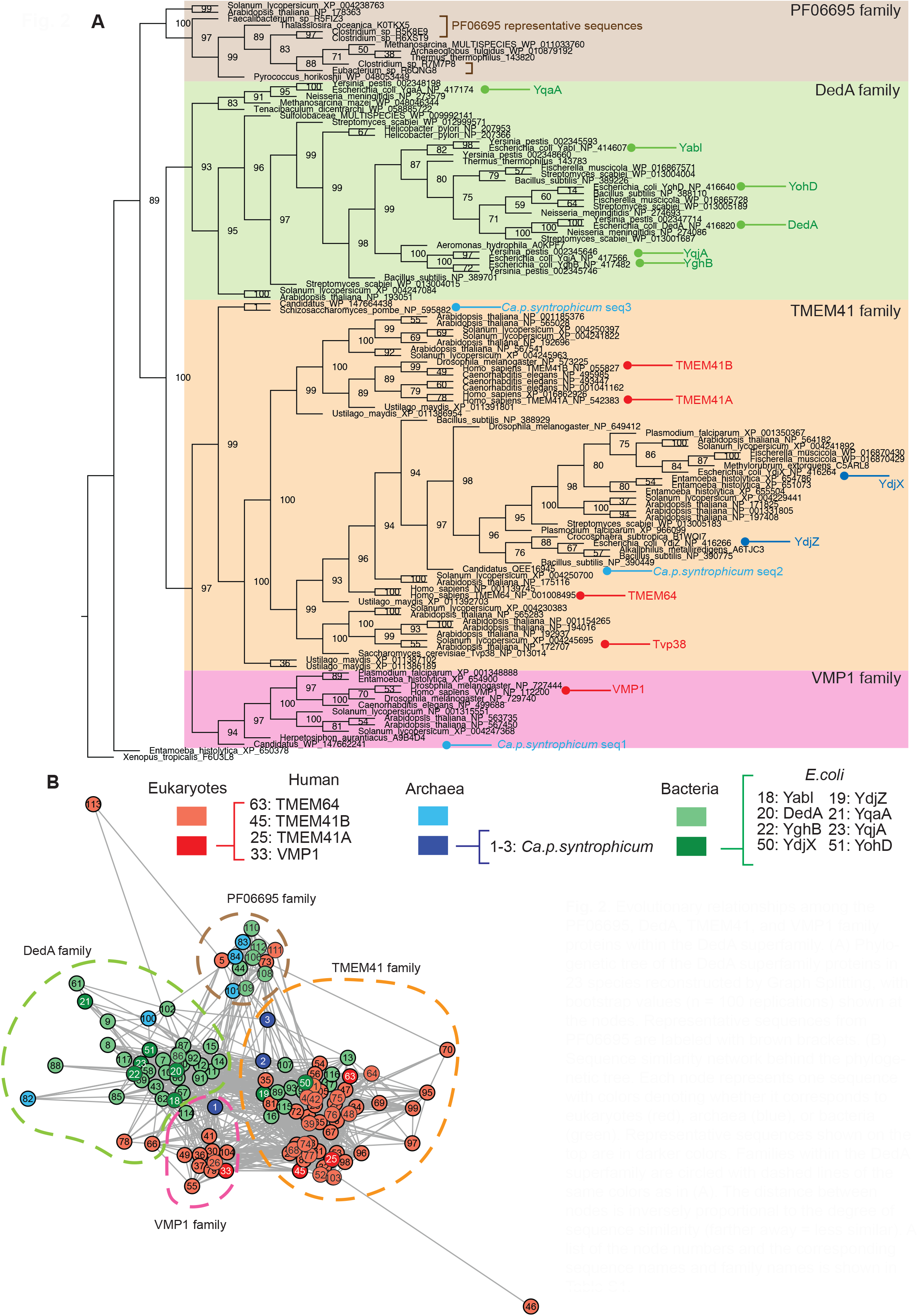
Evolutionary relationships among the PF06695, DedA, TMEM41, and VMP1 family proteins within the DedA superfamily. **(A)** Phylogenetic tree of the DedA superfamily proteins in 23 species reconstructed by Graph Splitting, with bootstrap values (*n* = 100 replications) shown at the nodes. Representative sequences from PF06695 are labeled with brown brackets. **(B)** Sequence similarity network behind the phylogenetic tree. Each node represents one sequence with colors denoting whether it corresponds to eukaryotes (red), archaea (blue), or bacteria (green). Representative sequences shown on the top are in darker colors. Families within the DedA superfamily are circled with dashed lines of the same colors as in (A). The distance between nodes is inversely proportional to the degree of sequence similarity (farther away = less similar). A list of the node numbers and the corresponding sequence names and family names is shown in Table S1.

### TMEM41A, TMEM64, and Tvp38 are not required for autophagy

The human genome encodes four DedA superfamily proteins: three TMEM41 family proteins (TMEM41A, TMEM41B, and TMEM64) and VMP1, whereas the genome of *Saccharomyces cerevisiae* encodes only one DedA superfamily protein, Tvp38, belonging to the TMEM41 family. Among these proteins, VMP1 and TMEM41B are known to be required for autophagy (Moretti et al., 2018; Morita et al., 2018; Ropolo et al., 2007; Shoemaker et al., 2019; Zhao et al., 2017). To determine whether the other three proteins are required for autophagy, we generated *TMEM41A* and *TMEM64* knockout (KO) HeLa cells and obtained a *tvp38*Δ yeast strain (BY4741). In wild-type (WT) unstarved HeLa cells, the amount of the autophagosome-localizing phosphatidylethanolamine-conjugated LC3 (LC3-II) increased upon treatment with bafilomycin A_1_, an inhibitor of vacuolar ATPase, indicating that autophagosomal LC3 was degraded in lysosomes by basal autophagy (Fig. 3A). Under starvation conditions, further accumulation of LC3-II was observed upon bafilomycin A_1_ treatment, suggesting an increase in autophagic flux during starvation. By contrast, in VMP1-KO cells, LC3-II accumulated even under nutrient-rich conditions, and the accumulation was not further increased by starvation or bafilomycin A_1_ treatment, suggesting that autophagic flux was blocked. Consistently, p62 and its phosphorylated form, which are selective substrates of autophagy, accumulated in VMP1-KO cells. Similarly, in TMEM41B-KO cells, LC3-II and phosphorylated p62 accumulated under both nutrient-rich and starvation conditions compared with WT cells, suggesting that autophagic activity was defective, although less severely than in VMP1-KO cells. Autophagic flux in TMEM41A-KO cells was normal as was previously shown in TMEM41A-knockdown cells (Morita et al., 2018) (Fig. 3A). TMEM64-KO cells also demonstrated normal autophagic flux.

**Fig. 3.**
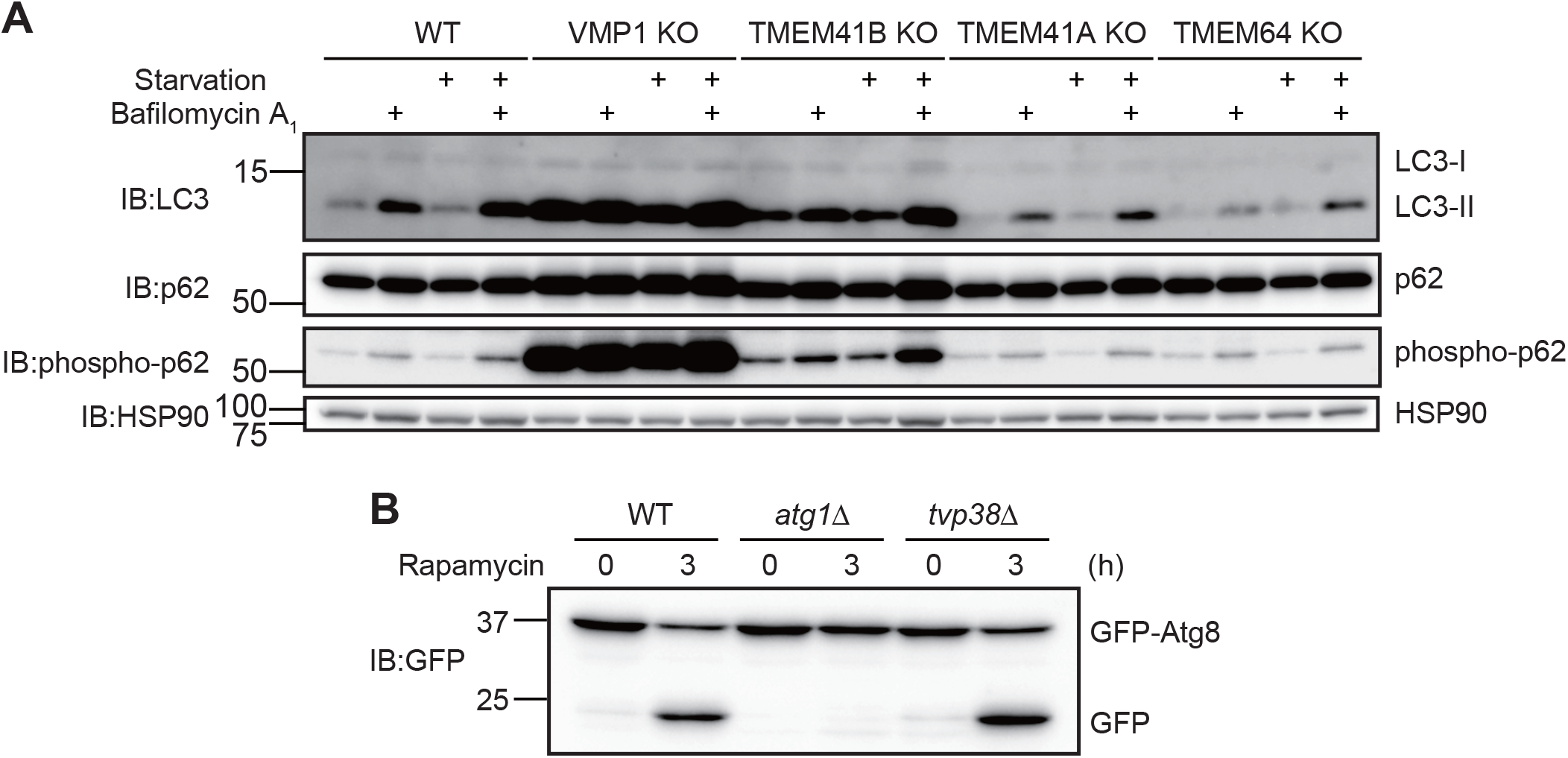
Autophagic activity in TMEM41A-KO, TMEM64-KO, and *tvp38*Δ cells. Autophagic flux of TMEM41A-KO and TMEM64-KO HeLa cells. Cells were cultured under nutrient-rich or starvation conditions with or without bafilomycin A_1_ for two hours. Data are representative of two independent experiments. **(B)** The GFP-Atg8 cleavage assay of wild-type, *atg1*Δ, and *tvp38*Δ. Cells were cultured with or without rapamycin for three hours. Data are representative of two independent experiments.

Autophagic flux in yeast was determined by monitoring the cleavage of GFP-Atg8, which was expressed in the cytosol and degraded after delivery to the vacuole by autophagy (Klionsky et al., 2016). In WT *Saccharomyces cerevisiae* (BY4741), cleaved GFP accumulated upon treatment with autophagy-inducible rapamycin, an inhibitor of TORC1. By contrast, in *atg1*Δ cells, cleaved GFP did not accumulate even after rapamycin treatment, indicating that autophagic activity was deficient. In *tvp38*Δ cells, cleaved GFP normally accumulated after rapamycin treatment, suggesting that autophagic activity was maintained (Fig. 3B). Thus, TMEM41A, TMEM64, and Tvp38 are not required for autophagy.

### Structural prediction of the DedA domain

Structural information of the DedA superfamily has been limited to secondary structure predictions, which suggested that members might contain 5–8 TMHs (Morita et al., 2018). However, the assignment of TMHs may not be accurate because even the conserved DedA domain is suggested to carry different numbers of TMHs depending on the species and proteins. To gain more reliable structural information and functional insights, we conducted *ab initio* structural prediction with trRosetta (Yang et al., 2020). Building on the assumption that coevolving residues are often in contact, trRosetta predicts distance and orientation between residues from sequence coevolution using deep learning. The accuracy of trRosetta prediction relies on the number and depth of the homologous sequences collected. In our case, 65535 homologous sequences (the default upper limit) were used for TMEM41A, TMEM41B, TMEM64 and YdjX, and 24330 homologous sequences were used for YdjZ (including overlapping sequences between them), yielding reliable structural predictions. On the other hand, predictions for VMP1 yielded results of low or medium quality owing to a relatively small number of homologous sequences. We therefore focused on the TMEM41 family in further analyses.

The prediction for TMEM41B generated a distance map with two ring-like patterns in the N- and C-terminal regions (Fig. 4A). Each of these ring-like patterns translates into a reentrant loop that enters the lipid bilayer but turns inside the membrane to exit from the same side. Notably, the first third of each ring was predicted to have contact (i.e. predicted distances less than 8 Å) between the two halves of the reentrant loops (e.g., L124–L135, Y121–L135, and Y117–S139 in reentrant loop 1 and L204–I215 and I201–S219 in reentrant loop 2) (Fig. 4B). Furthermore, the reentrant loops contain helix-breaking proline and glycine residues between the halves. Reentrant loop 1 turns roughly at the conserved proline-glycine residues (P130 and G131 in TMEM41B), and reentrant loop 2 turns at two conserved prolines separated by one or two other residues (P208 and P211 in TMEM41B) (Fig. 1; Fig. 4C). In addition to the contact areas within the reentrant loops, the contact map also suggests interactions between each of the reentrant loops and the TMHs, as well as between the TMHs, for example, contacts between the first half of reentrant loop 1 and TMH1 (the pink rectangle in Fig. 4A) and contacts between TMH1 and TMH2 (the orange rectangle in Fig. 4A). As shown, the two hairpin-shaped reentrant loops and two additional TMHs together form into a compact fold, suggesting that the DedA domain could be an independent structural domain.

**Fig. 4.**
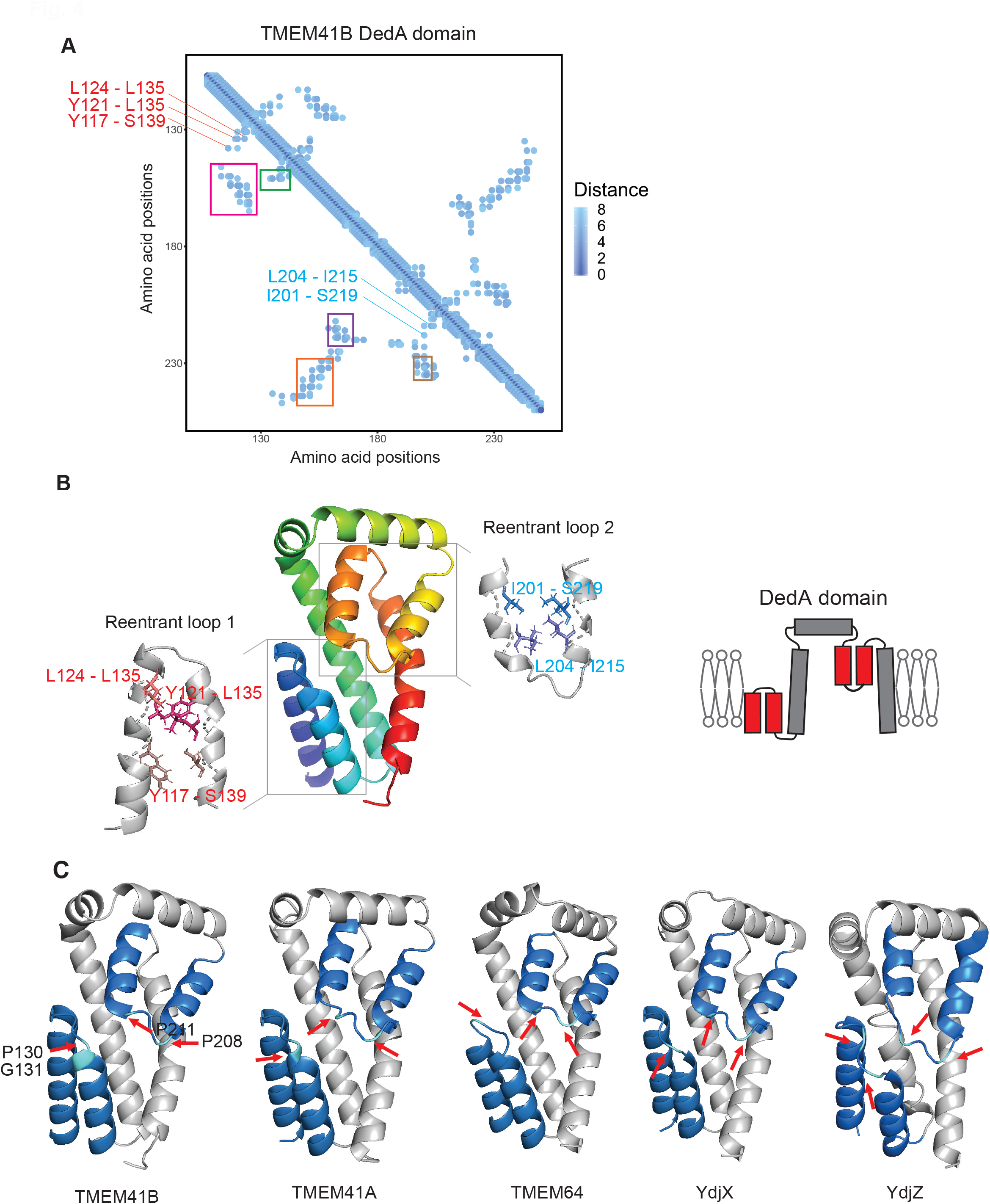
*Ab initio* structure prediction of the DedA superfamily proteins by trRosetta. **(A)** Distance map of the DedA domain of TMEM41B. The *x*- and *y*-axes show amino acid positions in TMEM41B, and the color gradient indicates the predicted distances between residue pairs. Examples of predicted interactions between the two halves of reentrant loops 1 and 2 are labeled in red and blue, respectively. The rectangles indicate contacts between the first half of reentrant loop 1 and TMH1 (pink), between the second half of reentrant loop 1 and TMH1 (green), between the second half of reentrant loop 2 and TMH1 (purple), between TMH1 and TMH2 (orange), and between the first half of reentrant loop 2 and TMH2 (brown). **(B)** Top-ranking model of TMEM41B predicted by trRosetta, with reentrant loops 1 and 2 enlarged. Different colors indicate pairs of residues predicted to be in contact (i.e. with predicted distance less than 8 Å) by coevolution. A model of membrane topology of the DedA domain is also shown. **(C)** Predicted models of TMEM41B, TMEM41A, TMEM64, YdjX, and YdjZ, with reentrant loops in dark blue and conserved proline residues in light blue. Conserved proline residues at which the reentrant loops turn are labeled with red arrows.

The predictions for TMEM41A, TMEM64, YdjX, and YdjZ all yielded similar contact maps (Fig. S2) and structures (Fig. 4C). Along this line, we found that the GREMLIN structural prediction server, which also uses coevolution information, lists similar contact maps and structures for R6BJC6 protein representing the PF06695 family (Ovchinnikov et al., 2014). We were also able to obtain similar contact maps and predicted structures using a different prediction method EVfold (Hopf et al., 2019) (Fig. S3).

The predicted structure visualizes a characteristic structure of two reentrant loops facing each other in the membrane. In order to gain insights into its molecular function, we searched the PDBTM database (Kozma et al., 2013), a database of annotated transmembrane proteins with solved structures, for other proteins with two reentrant loops (keyword “0 [type] AND 2 [n_loop]” in advanced search). Such reentrant loops were found in transporters and ion channels such as aquaporins (AQPs), chloride channels (CLCs), solute carrier family 1 (SLC1), solute carrier family 13 (SLC13), solute carrier family 28 (SLC28), and bacterial undecaprenyl pyrophosphate phosphatase (UppP) (Chang et al., 2014; Forrest, 2015; Kanai et al., 2013; Screpanti and Hunte, 2007) (Fig. S4A; Table S2). Among them, the topology of the substrate-binding region of SLC1 is most similar to that of the DedA domain: both consist of two repeats of a reentrant loop and a succeeding TMH, and the membrane topologies of the two repeats are inverted (Kanai et al., 2013). Consistently, SLC1 shows a coevolution pattern similar to that of the DedA domain (Fig. S4B,C). Thus, the two facing reentrant loops of the DedA domain might also serve as a substrate-binding site for potential ion-coupled transporters.

### Biochemical verification of the topology of TMEM41B

Structural prediction by trRosetta and EVfold suggests that the DedA domain contains reentrant loop 1, TMH1, an extra-membrane region, reentrant loop 2, and TMH2 (from the N to C terminus) (Fig. 4). In addition, TMHMM predicted that TMEM41B has two more TMHs outside of the DedA domain, at the N- and C-terminal ends (Fig. 5A; Fig. S4A) (Moller et al., 2001). To verify the predicted topology of TMEM41B experimentally, we performed substituted cysteine accessibility method (SCAM) analysis (Bogdanov et al., 2005). Cysteine has a thiol group that can be conjugated with maleimide or maleimide-containing molecules such as methoxypolyethylene glycol maleimide (PEG-maleimide) and *N*-ethylmaleimide (NEM). Once a protein is conjugated with PEG-maleimide, it becomes larger and can be separated from an unconjugated form by SDS-PAGE (Fig. 5B) (Davis et al., 2019). As PEG-maleimide is cell-impermeable, specific labeling is achieved only after membrane permeabilization with a detergent (Fig. 5C).

**Fig. 5.**
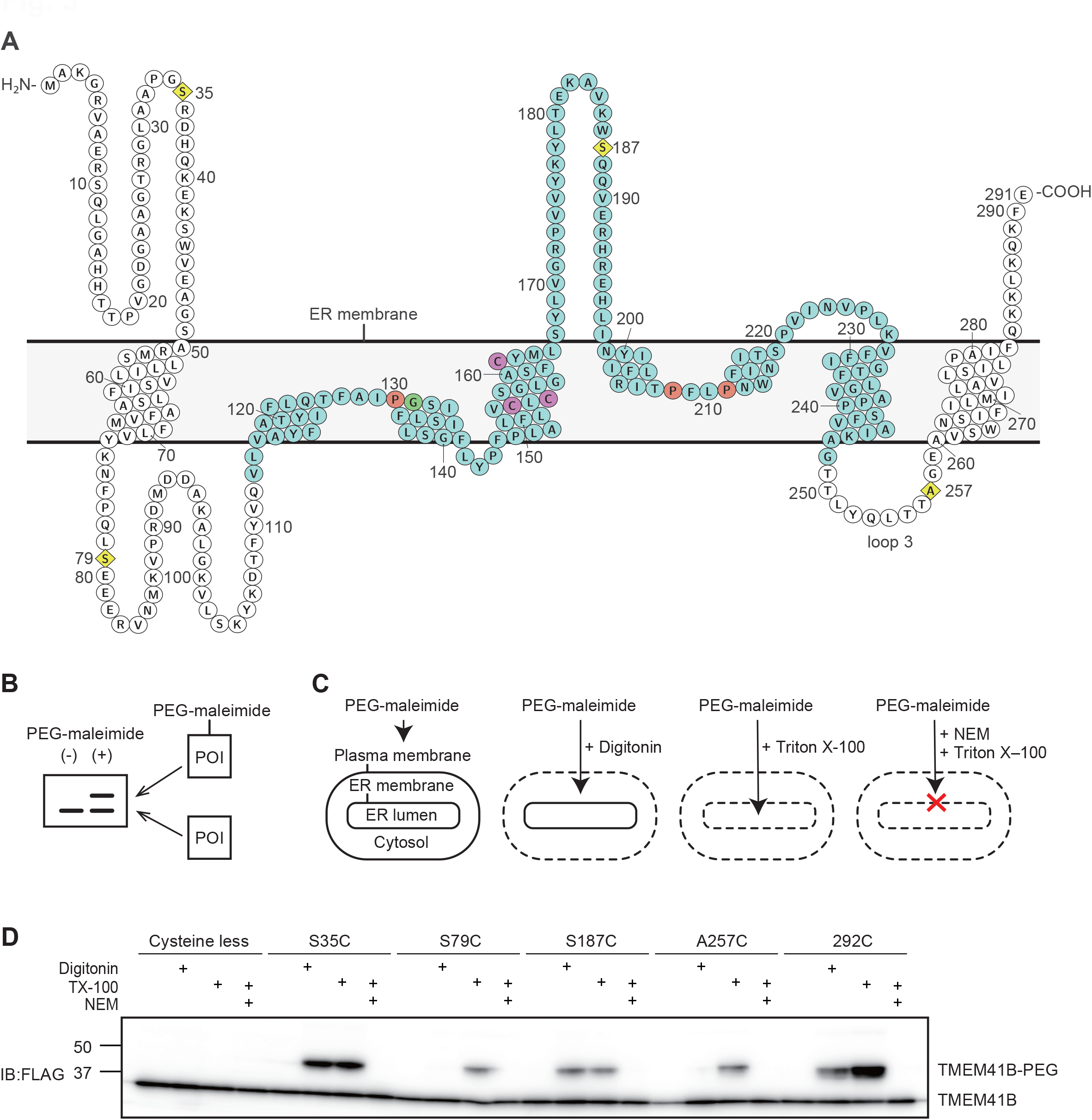
SCAM-based topology analysis of TMEM41B. **(A)** Residues in the DedA domain are in blue circles. Residues mutated in the SCAM analysis (yellow diamond), conserved prolines (orange circle) and glycines (green circle) located in the reentrant loops, and endogenous cysteines (magenta circle) are marked. Schematic representation of the effect of PEG-maleimide treatment in SDS-PAGE. If PEG-maleimide is conjugated with a protein-of-interest (POI), a higher molecular weight band will appear. **(C)** Experimental design for protein topological analysis of ER proteins. Dotted lines indicate permeabilized membranes. When the plasma membrane is permeabilized by digitonin, PEG-maleimide can be conjugated with cysteine residues located in the cytosol. When the ER membrane is permeabilized by Triton X-100, PEG-maleimide is able to penetrate into the ER lumen and be conjugated with cysteine residues there. Conjugation between PEG-maleimide and cysteine is inhibited by NEM treatment. **(D)** SCAM-based topology analysis of TMEM41B. HeLa cells expressing each mutant were treated with indicated reagents. Mutants were constructed on a cysteine-less TMEM41B background. The 292C mutant is cysteine-less TMEM41B with an additional cysteine on the C terminus. Data are representative of two independent experiments.

We prepared cells expressing cysteine-less TMEM41B or its variants in which one of the amino acids was replaced with cysteine (yellow residues in Fig. 5A). Upon PEG-maleimide (molecular weight = 5,000) treatment, cysteine-less TMEM41B did not show any band shift, even in the presence of the mild detergent digitonin (Fig. 5D). By contrast, the single-cysteine TMEM41B mutants S35C, S187C, and 292C (in which cysteine was added to the C terminus) showed an additional high-molecular-weight band in the presence of both PEG-maleimide and digitonin, which permeabilized the plasma membrane. The intensity of these high-molecular-weight bands was unchanged upon Triton X-100 treatment, which also permeabilizes organellar membranes (Fig. 5C, D). Formation of these bands was inhibited by pretreatment with NEM, which blocked the conjugation between PEG-maleimide and the thiol group of cysteine. These results suggest that these high-molecular-weight bands represent PEG-conjugated TMEM41B and that S35, S187, and the C terminus are in the cytosol. On the other hand, the single-cysteine mutants S79C and A257C did not produce TMEM41B-PEG bands in the presence of digitonin, but did so in the presence of Triton X-100, suggesting that these residues are present in the lumen of the ER (Fig. 5D).

We also tested whether these single-cysteine TMEM41B mutants retained their original topologies by assessing their function in autophagy. In TMEM41B KO cells, the amount of LC3-II accumulated, representing a block in autophagic flux (Moretti et al., 2018; Morita et al., 2018; Shoemaker et al., 2019). This defect was restored by exogenous expression of WT TMEM41B or all the single-cysteine TMEM41B mutants, suggesting that these mutants are correctly integrated into the membrane (Fig. S5). Thus, these results verified the topology predicted by trRosetta and EVfold; the presence of S79 and S187 on opposite sides of the membrane suggests that reentrant loop 1 indeed turns back in the membrane rather than penetrating the membrane, and the presence of S187 and A257 on opposite sides suggests that reentrant loop 2 is indeed reentrant. Collectively, these results suggest that TMEM41B is composed of four TMHs and two reentrant loops facing each other, and both the N and C terminus are in the cytosol.

## DISCUSSION

### Evolution of the DedA superfamily proteins and acquisition of autophagic function

In this study, we conducted a phylogenetic analysis of the DedA superfamily, containing the DedA, TMEM41, VMP1, and PF06695 families, all of which possess the DedA domain. While the DedA and PF06695 families and the TMEM41 and VMP1 families primarily contain prokaryotic and eukaryotic proteins, respectively, each of these four families contains both prokaryotic and eukaryotic proteins (Fig. 2), indicating the origins of these four families predate the split between prokaryotes and eukaryotes. Among the DedA superfamily members, only VMP1 and TMEM41B have a role in autophagosome formation, whereas TMEM41A, TMEM64, and Tvp38 do not (Fig. 3) (Moretti et al., 2018; Morita et al., 2018; Shoemaker et al., 2019). Notably, although prokaryotes do not have an autophagy system or lysosomes, they do have ancestors of both VMP1 and TMEM41. There are two possible scenarios of how the prokaryotic DedA ancestors acquired autophagic functions in the course of evolution. One is that the prokaryotic ancestors of VMP1 and TMEM41 had a common function at the plasma membrane, which was later directly used in autophagy in eukaryotes. In this case, these proteins might have spontaneously acquired their autophagic function after translocation to the ER membrane. However, this hypothesis cannot explain why most TMEM41 family members do not have autophagic function. Even TMEM41A, the closest homolog of TMEM41B, does not play a role in autophagy. Also, Tvp38, the only TMEM41 family protein in yeast, is dispensable for autophagy.

The alternative scenario is that a VMP1 ancestor acquired autophagic function during evolution first, probably in a eukaryotic ancestor. Accordingly, the autophagic function of VMP1 is conserved broadly in eukaryotes, such as in Metazoa and Amoebozoa (Calvo-Garrido et al., 2008) and likely in green algae (Tenenboim et al., 2014). Later, probably after diverging from TMEM41A, TMEM41B became involved in autophagy, and this new function of TMEM41B occurred dependently on the preexisting autophagic function of VMP1, for example, through binding to VMP1 (Morita et al., 2018). This hypothesis can explain why only TMEM41B is involved in autophagy among the TMEM41 family proteins. It is also consistent with the previous observation that the role of TMEM41B is rather accessory; the phenotype of TMEM41B-KO cells is milder than that of VMP1-KO cells, and overexpression of VMP1 can rescue the phenotype of TMEM41B-KO cells, but the opposite cannot (Moretti et al., 2018; Morita et al., 2018; Shoemaker et al., 2019). A more comprehensive analysis of the function of VMP1 and TMEM41 family proteins in non-metazoan eukaryotes will further provide crucial information on how DedA family proteins acquired their autophagic function.

### Potential functions of the DedA superfamily proteins based on the predicted structure

The next fundamental unresolved issue is the function of the evolutionarily conserved DedA superfamily proteins. Starting with the HMM-based alignment, we first specified the core domain conserved in these proteins, which was rather ambiguously specified before (Doerrler et al., 2013; Keller and Schneider, 2013; Morita et al., 2018), and we defined it as the DedA domain. Our predicted structure of the DedA domain is unique; it contains two reentrant loops and two TMHs. This topology was verified experimentally (Fig. 5). Thus, the structure of the proposed DedA domain differs from the previous speculation that the DedA family proteins adopt half of a LeuT fold-like structure (Keller et al., 2014; Khafizov et al., 2010). During the preparation of this manuscript, a similar structural prediction was reported in bioRxiv (Mesdaghi et al., 2020).

The reentrant loops in transporters and ion channels often directly interact with substrates (Johnson et al., 2012; Kanai et al., 2013; Mancusso et al., 2012; Tornroth-Horsefield et al., 2010; Workman et al., 2018). Among them, the local architecture around the substrate-binding site of the Na^+^-coupled glutamate transporter SLC1 (Fig. S4A) is highly similar to that of the DedA domain (Kanai et al., 2013). Therefore, the DedA domain might have an ion-coupled transport function. It is tempting to speculate that VMP1 and TMEM41B are Ca^2+^-coupled transporters because VMP1 physically interacts with and is functionally related to SERCA, a Ca^2+^ transporter in the ER (Zhao et al., 2017). With regard to the potential substrate, we note that bacterial UppP uses a similar pair of reentrant loops to bind the head groups of membrane lipids and catalyze their dephosphorylation within the membrane (Workman et al., 2018). Although the catalytic residues are not conserved in the DedA domain and the overall topologies are not identical between UppP and the DedA domain (additional elements, including two TMHs, are inserted between the two internal repeats in UppP), the linkage between this structural feature and the lipid binding of UppP suggests that the DedA domain may recognize the head groups of membrane lipids as substrates. This hypothesis aligns well with the lipid-related phenotypes observed in VMP1- and TMEM41B-deficient eukaryotic cells (Calvo-Garrido et al., 2008; Moretti et al., 2018; Morishita et al., 2019; Morita et al., 2018; Ropolo et al., 2007; Shoemaker et al., 2019; Zhao et al., 2017), and YqjA- and YghB-deficient bacterial cells (Boughner and Doerrler, 2012; Thompkins et al., 2008). The slight phenotypic difference between VMP1 and TMEM41B deficiency may represent a difference in their substrates. Furthermore, several genetic studies suggest ion-dependent solute exporting functions for YqjA and YghB (Boughner and Doerrler, 2012; Doerrler et al., 2013; Keller et al., 2015; Kumar et al., 2016; Kumar and Doerrler, 2014; Ledgham et al., 2005; Panta et al., 2019). Collectively, predicted structural similarities suggest that the DedA superfamily proteins could have ion-dependent lipid or solute transport functions.

Reentrant loops are generally not hydrophobic enough to be stably embedded in membranes; they are often stabilized by surrounding TMHs (Yan and Luo, 2010) and/or participate in the subunit interface in a complex (Table S2). Thus, DedA superfamily proteins may also form similar complexes. Determining the actual structure of the DedA superfamily will eventually reveal the function of this broadly conserved family of proteins.

## MATERIALS AND METHODS

### Remote homology search

Remote homology search was conducted using HHsearch (Steinegger et al., 2019), part of the MPI Bioinformatics Toolkit (Zimmermann et al., 2018). The “local:realign” option was used to enable the maximum accuracy algorithm for more accurate alignment. Full-length VMP1 sequence (NP112200.2) was used as the query, and Pfam (El-Gebali et al., 2019) and the proteomes of *Homo sapiens, Saccharomyces cerevisiae, Escherichia coli*, and twenty randomly selected eukaryotic, bacterial, and archaeal species (Fig. S1) were used as the search database. A total of 125 homologous sequences (E-value cutoff = 1), including one representative sequence from PF09335 SNARE associated Golgi protein and PF06695 putative small multi-drug export protein each, were identified. PF09335 was renamed as the VTT family (Morita et al., 2018). The alignment in Fig. 1 was generated by MUSCLE v3.8.1551 (Edgar, 2004) together with 500 randomly selected sequences from a pool of TMEM41B- and VMP1-homologous sequences and PF06695 representative sequences.

### Phylogeny reconstruction

The phylogenetic tree of the DedA superfamily was reconstructed using Graph Splitting (Matsui and Iwasaki, 2020) with default parameters. In addition to the homologous sequences identified, five additional sequences from PF09335 and PF06695 each as well as three sequences from the archaeon *Ca. P. syntrophicum* were also included. After eliminating redundant sequences with 95% sequence similarity, 117 sequences remained. Randomly selecting different sets of sequences from these two Pfam families did not change the result. While Graph Splitting does not estimate branch length, bootstrap values (number of replicates = 100) are displayed at the nodes of the phylogenetic tree.

### Structural prediction based on coevolution

Structural prediction was conducted by trRosetta (Yang et al., 2020) and EVfold (Hopf et al., 2019) with default parameters. For trRosetta, residue pairs with predicted distance less than 8 Å were used to make the distance plot. As the five predicted models produced for each protein were highly similar, the first model was plotted. For EVfold, at the multiple-sequence-alignment-building step, significance thresholds of 0.2, 0.3, and 0.4 (bitscore/sequence length) were tested, and 0.4 was ultimately chosen because it produces the clearest ring-like pattern on the N-terminal side (Fig. S3). For each protein, the top-ranking model was selected. The models were plotted using Pymol.

### Antibodies

The following antibodies were used for immunoblotting: mouse monoclonal antibodies against HSP90 (610419; BD), FLAG (F1804; Sigma-Aldrich), and GFP (11814460001; Roche) and rabbit polyclonal antibodies against p62/SQS TM1 (PM045; MBL) and phospho-p62 (PM074; MBL). Rabbit polyclonal antibody against LC3 was described previously (Hosokawa et al., 2006). Peroxidase-conjugated anti-mouse and anti-rabbit immunoglobulins (111-035-144; Jackson ImmunoResearch Laboratories, Inc.) were used as secondary antibodies.

### Plasmid

The pMRXIP-TMEM41B-3xFLAG encoding human TMEM41B was described previously (Morita et al., 2018) and used as a template for making the mutants. Three endogenous cysteines at 153, 155, and 163 were mutated to serines, and the product was utilized to make each single cysteine mutant. PrimeSTAR Max DNA Polymerase (R045A; Takara Bio Inc.) was used for mutagenesis. Preparations of primers and mutagenesis steps followed the manufacturer’s instructions. Each generated construct was confirmed by sequencing (Eurofins Genomics JP). For generation of knockout cell lines, guide RNA (gRNA) targeting TMEM41A (5’-GCCGAGAAGCGGGCGCATGT-3’) and TMEM64 (5’-CCGCGCTGGGCCGAGGCATG-3’) were cloned into pSpCas9(BB)-2A-GFP (Addgene #48138; a gift from Dr. F. Zhang, Broad Institute of Massachusetts Institute of Technology and Harvard, USA). Additionally, pRS316-GFP-Atg8 was used for GFP-Atg8 assay.

### Cell culture

Using Dulbecco’s modified Eagle’s medium (DMEM [D6546; Sigma-Aldrich]) supplemented with 10% fetal-bovine-serum (FBS) and 2 mM glutamine (25030-081; Gibco), cells were cultured in a 5% CO_2_ incubator. To impose starvation conditions, cells were washed with phosphate-buffered saline (PBS) and cultured in amino acid-free DMEM (048-33575; Wako) without FBS. For vacuolar ATPase inhibition, cells were cultured with 100 nM bafilomycin A_1_ (B1793; Sigma-Aldrich) for 2 hours.

### Generation of stable cell lines

HEK293T cells were transiently transfected with a target plasmid, pCG-VSV-G, and pCG-gag-pol (gifts from Dr. T. Yasui, Osaka University, Japan) using Lipofectamine 2000 (11668019; Thermo Fisher Scientific). Two days after transfection, culture medium including retrovirus was collected through a 0.45-μm syringe filter unit (SLHV033RB; Merck Millipore). Retrovirus was mixed with 8 µg/mL polybrene (H9268; Sigma-Aldrich) and host cells were transfected with it. After 24 hours, the medium was exchanged to DMEM containing 2 µg/mL puromycin (P8833; Sigma-Aldrich) for selection.

### Establishment of *TMEM41A*-KO and *TMEM64*-KO HeLa cells

HeLa cells were transfected with pSpCas9(BB)-2A-GFP encoding gRNAs using FuGENE HD Transfection Reagent (E2311; Promega). Two days after transfection, GFP-positive cells were isolated by cell sorter (MoFlo Astrios EQ; Beckman Coulter), and single clones were obtained. Clones containing knockout mutations were selected by immunoblotting and sequencing of genomic DNA.

### Immunoblotting

Cells were collected in ice cold PBS with a cell scraper and centrifuged at 5000 × *g* for 3 minutes. They were then treated with lysis buffer (1% Triton X-100, 50mM Tris-HCl pH 7.5, 150 mM NaCl, 1mM EDTA, protease inhibitor cocktail [03969; Nacalai Tesque]) and incubated on ice for 15 minutes. Then, lysed samples were centrifuged at 12,000 × *g* for 15 minutes, and the resulting supernatants were collected for analysis. SDS-PAGE sample buffer (46.7 mM Tris-HCl, pH 6.8, 5% glycerol, 1.67% sodium dodecyl sulfate, 1.55% dithiothreitol, and 0.02% bromophenol blue) was added to samples and boiled. The SDS-PAGE was conducted to separate proteins, which were transferred to a polyvinylidene difluoride (PVDF) membrane. Appropriates antibodies were applied to the membrane after blocking with Tris-buffered saline with Tween 20 (TBST) containing 5% skim milk. Membranes were incubated with primary antibodies at 4°C overnight, followed by incubation with secondary antibodies at room temperature for an hour. After washing and reacting the membranes with Super-Signal West Pico Chemiluminescent substrate (1856135; Thermo Fisher Scientific), signals were detected by FUSION Solo S (Vilber-Lourmat). Contrast and brightness adjustments were performed using Fiji software (Schindelin et al., 2012).

### Modification of cysteine residues

Cells were transiently transfected with the indicated mutants using Lipofectamine 2000 (11668019; Thermo Fisher Scientific). After 24 hours, the plasma membrane was permeabilized using DMEM containing 100 μg/mL digitonin (12333-51; Nacalai Tesque) for 3 minutes at 37°C. To permeabilize both the plasma and ER membranes, cells were treated with 0.1% Triton X-100 in PBS containing 0.1 mM CaCl_2_ and 1 mM MgCl_2_ (PBSCM) for 3 minutes at room temperature. Detergents were removed, and cells were washed with PBS. Then, *N*-ethylmaleimide (NEM [15512-11; Nacalai Tesque]) was diluted to 5.0 mM using PBS, and cells were incubated for 1 hour on ice using a rocker before permeabilization with 0.1% Triton X-100. Methoxypolyethylene glycol maleimide (PEG-maleimide [63187; SIGMA]) was diluted to 1.5 mM using PBSCM. After membrane permeabilization, cells were incubated in PEG-maleimide solutions for 30 minutes on ice using a rocker. PEG-maleimide modification was stopped by a solution containing 10 mM dithiothreitol (DTT [14112-52; Nacalai Tesque]) in PBSCM with 2% bovine serum albumin (BSA) for 10 minutes on ice using a rocker. After removing the solution, cells were collected in ice cold PBS and centrifuged at 5000 × *g* for 3 minutes. They were broken by passaging with a 26-gauge needle 20 times with lysis buffer (0.1% Triton X-100, 250 mM sucrose, 25 mM Tris-HCl (pH 7.5), 2 mM DTT, protease inhibitor cocktail [03969; Nacalai Tesque]) and incubated on ice for 15 minutes. Finally, lysates were centrifuged at 100 × *g* for 10 minutes, and supernatants were collected as samples.

### Yeast cells and GFP-Atg8 cleavage assay

The yeast knock out haploid MATa collection (TKY3502; TOT) was obtained from Funakoshi Co., Ltd. After confirmation of knockout by polymerase chain reaction (PCR), the *tvp38*Δ and *atg1*Δ strains were used for the experiment. Cells were transformed with pRS316-GFP-Atg8 as previously described (Gietz and Woods, 2002). A GFP-Atg8 cleavage assay was performed as previously described (Cheong and Klionsky, 2008).

## Acknowledgements

We would like to thank Drs. Wataru Iwasaki, Motomu Matsui, and Euki Yazaki for helpful discussions on the phylogenetic analysis conducted.

## Figure Legends

**Figure S1**. Phylogenetic tree of *Homo sapiens, Saccharomyces cerevisiae, Escherichia coli*, and twenty randomly selected species spanning eukaryotes, bacteria, and archaea included in the remote homology search.

**Figure S2**. Distance maps of the DedA domains of the TMEM41 family proteins inferred by trRosetta. The *x*- and *y*-axes represent amino acid positions in each protein, and the color gradient shows the predicted distances between residue pairs.

**Figure S3**. *Ab initio* structure prediction of the DedA superfamily proteins by EVfold. **(A**,**B**,**C)** Contact maps of the DedA domain in TMEM41B with bitscore/sequence length threshold = 0.2, 0.3 or 0.4. The x- and y-axes show amino acid positions, and the color gradient indicates the probability that residue pairs are evolutionarily coupled. **(D**,**E**,**F**,**G)** Contact maps of the DedA domain in TMEM41A, TMEM64, YdjX and YdjZ with bitscore/sequence length threshold = 0.4. **(H)** Top-ranking models with the same coloring scheme as Fig. 4C.

**Figure S4**. Structural similarity between *Pyrococcus horikoshii* glutamate transporter solute carrier family 1 (SLC1) and the DedA domain. **(A)** Membrane topologies of TMEM41B, SLC1, SLC13, SLC28, and undecaprenyl pyrophosphate phosphatase (UppP). Reentrant loops are indicated (red). **(B)** Distance map of SLC1 with the two reentrant loops labeled by red rectangles. **(C)** Structure of SLC1 (PDB: 1xfh) with the reentrant loops colored in pink. The proline and glycine residues at the turns are indicated by red arrows. The first two TMHs are omitted for clarity.

**Figure S5**. The single cysteine TMEM41B mutants retained autophagic function. TMEM41B-KO HeLa cells stably expressing cysteine-less TMEM41B (ΔCys) or the single cysteine mutants were starved with or without bafilomycin A_1_ treatment. Cell lysates were subjected to immunoblotting using anti-p62, anti-phospho-p62, and anti-LC3 antibodies.

**Table S1. DedA superfamily proteins analyzed in Fig. 2**. A list of the numbers and corresponding protein names in Fig. 2B. For each protein, the family it belongs to and whether it is eukaryotic, bacterial, or archaeal are also shown.

**Table S2. Transmembrane proteins containing 2 loops found in PDBTM**. The PDBTM database was searched for proteins with two reentrant loops using the keywords “0 [type] AND 2 [n_loop]”, and structures were checked one by one to investigate if the reentrant loops were facing each other.

