## Supplementary material for "Evolution and insights into the structure and function of the DedA superfamily containing TMEM41B and VMP1": Figure S1 - S5, Table S1 - S2

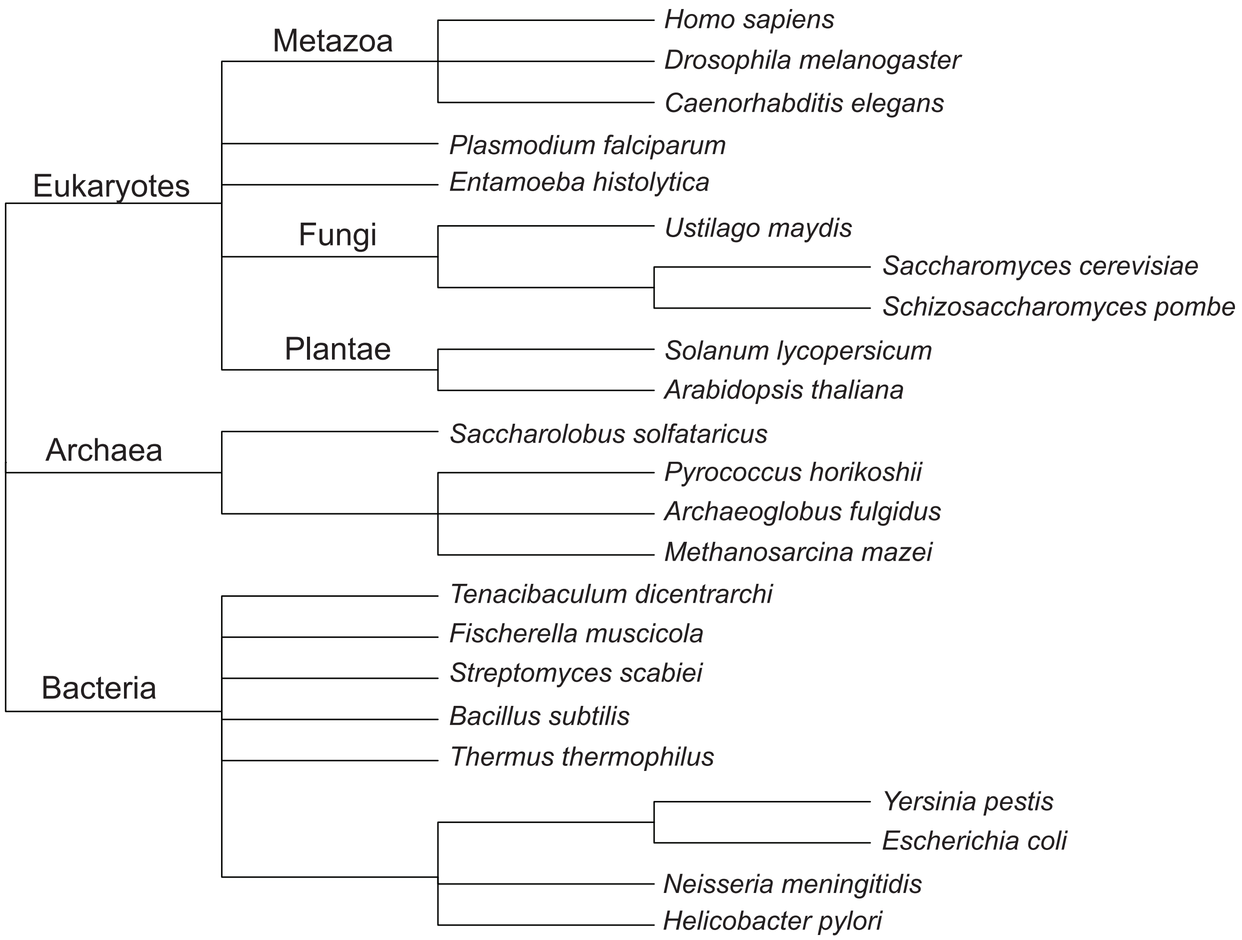

**Figure S1**. Phylogenetic tree of Homo sapiens, Saccharomyces cerevisiae, Escherichia coli, and twenty randomly selected species spanning eukaryotes, bacteria, and archaea included in the remote homology search.

**Figure S2**

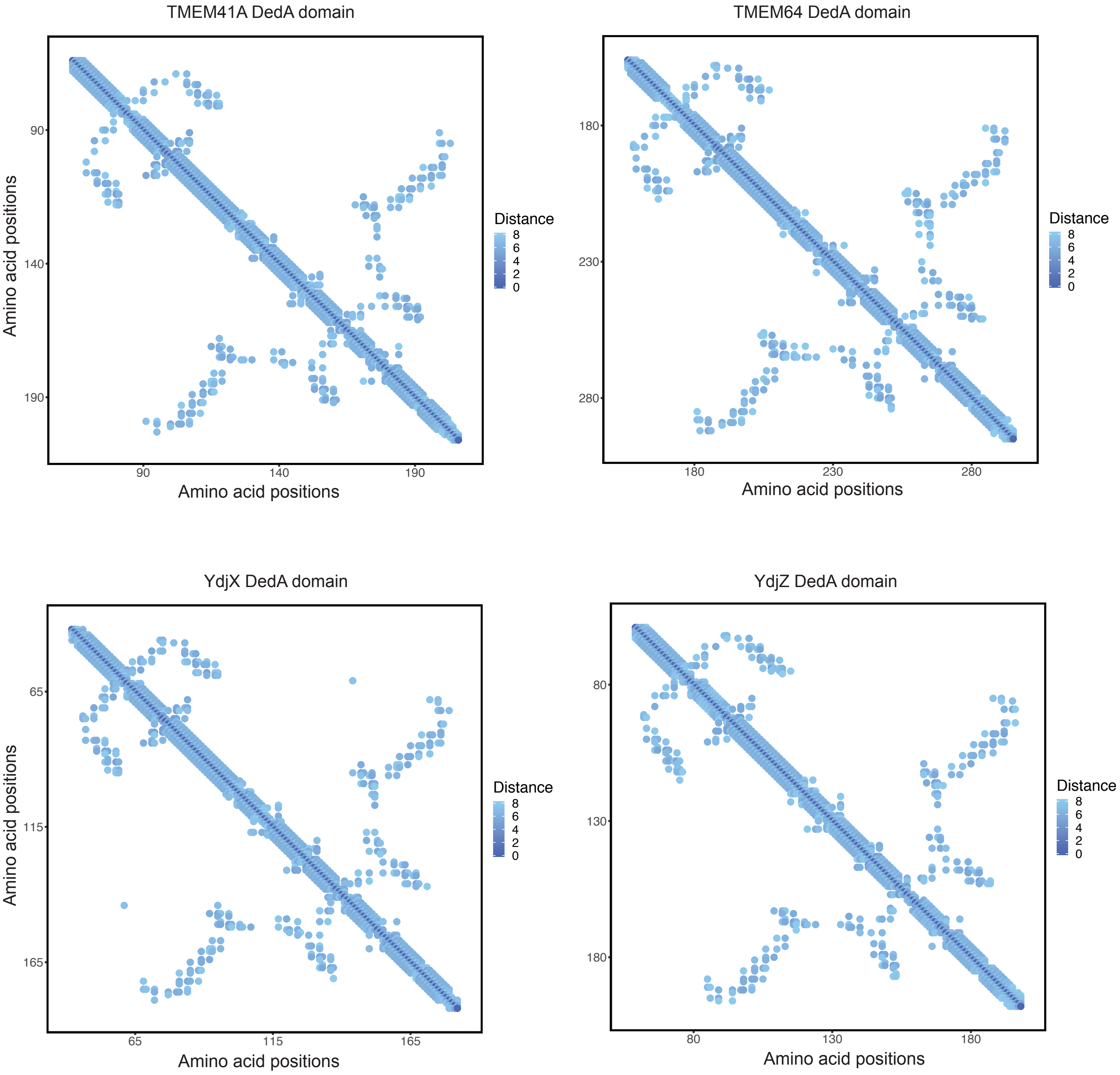

**Figure S2**. Distance maps of the DedA domains of the TMEM41 family proteins inferred by trRosetta. The x- and y-axes represent amino acid positions in each protein, and the color gradient shows the predicted distances between residue pairs.

### Figure S3

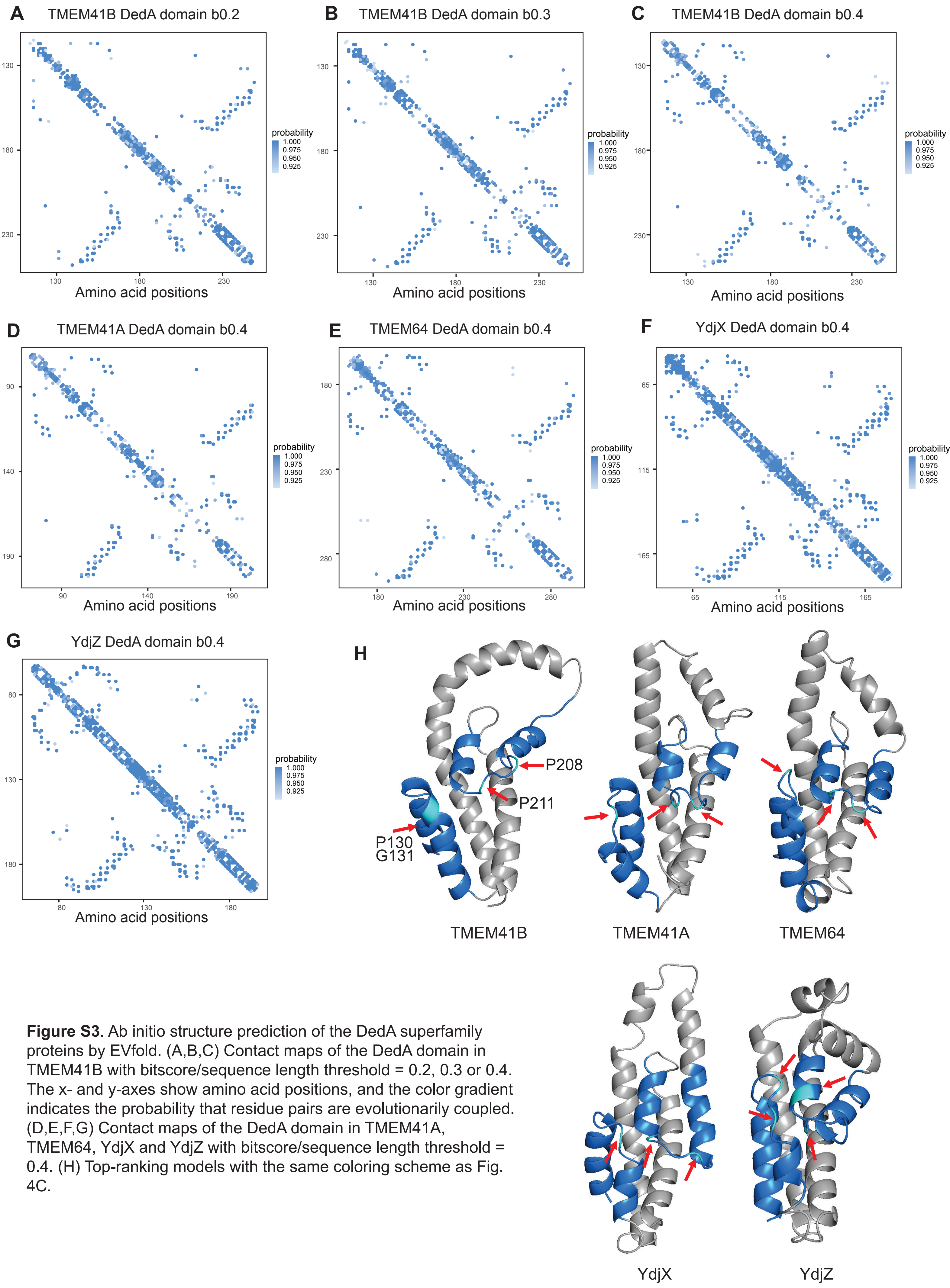

**Figure S3**. Ab initio structure prediction of the DedA superfamily proteins by EVfold. (A,B,C) Contact maps of the DedA domain in TMEM41B with bitscore/sequence length threshold = 0.2, 0.3 or 0.4. The x- and y-axes show amino acid positions, and the color gradient indicates the probability that residue pairs are evolutionarily coupled. (D,E,F,G) Contact maps of the DedA domain in TMEM41A, TMEM64, YdjX and YdjZ with bitscore/sequence length threshold = 0.4. (H) Top-ranking models with the same coloring scheme as Fig. 4C.

**Figure S4**

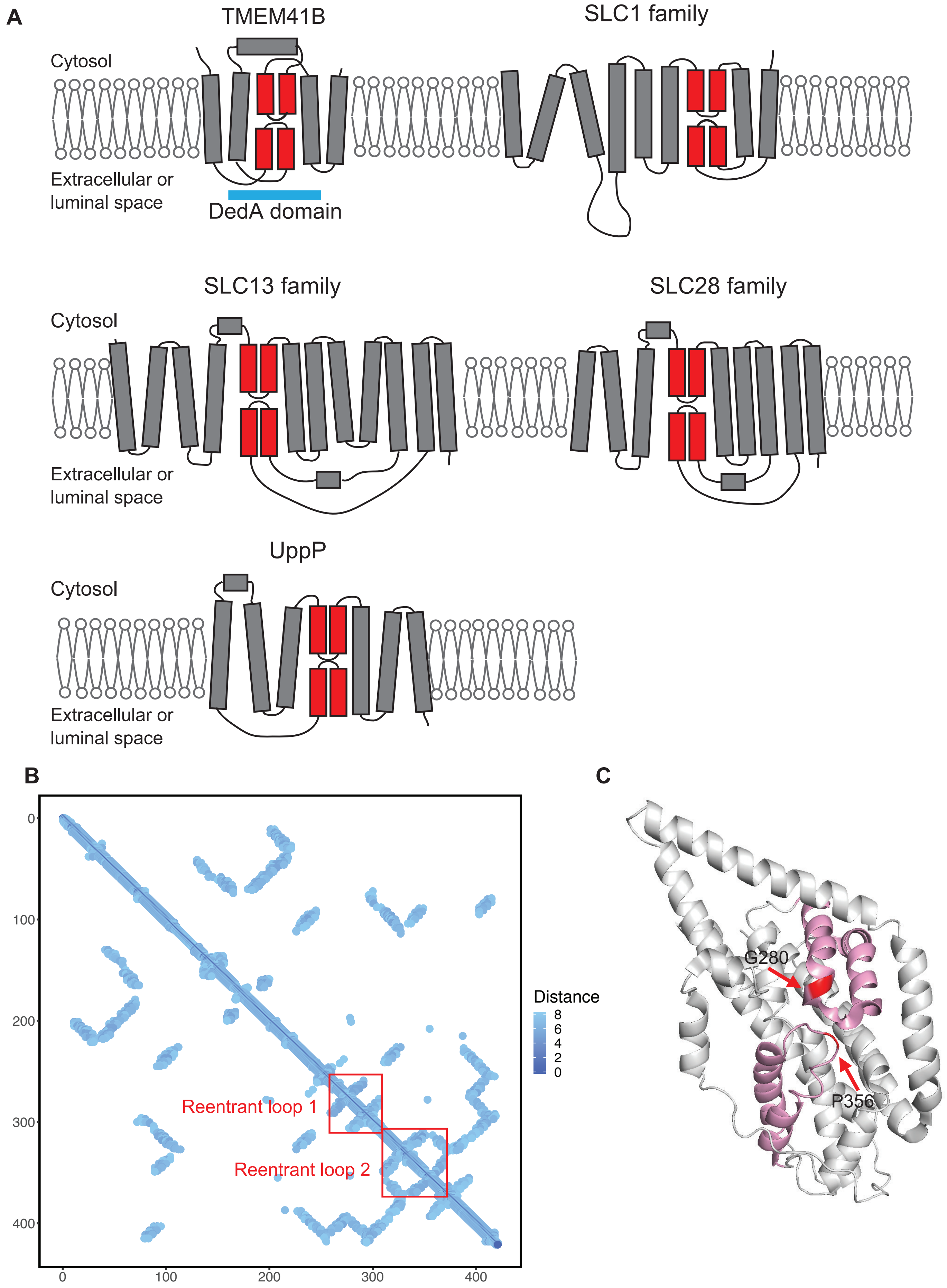

**Figure S4**. Structural similarity between *Pyrococcus horikoshii* glutamate transporter solute carrier family 1 (SLC1) and the DedA domain. (A) Membrane topologies of TMEM41B, SLC1, SLC13, SLC28, and undecaprenyl pyrophosphate phosphatase (UppP). Reentrant loops are indicated (red). (B) Distance map of SLC1 with the two reentrant loops labeled by red rectangles. (C) Structure of SLC1 (PDB: 1xfh) with the reentrant loops colored in pink. The proline and glycine residues at the turns are indicated by red arrows. The first two TMHs are omitted for clarity.

### Figure S5

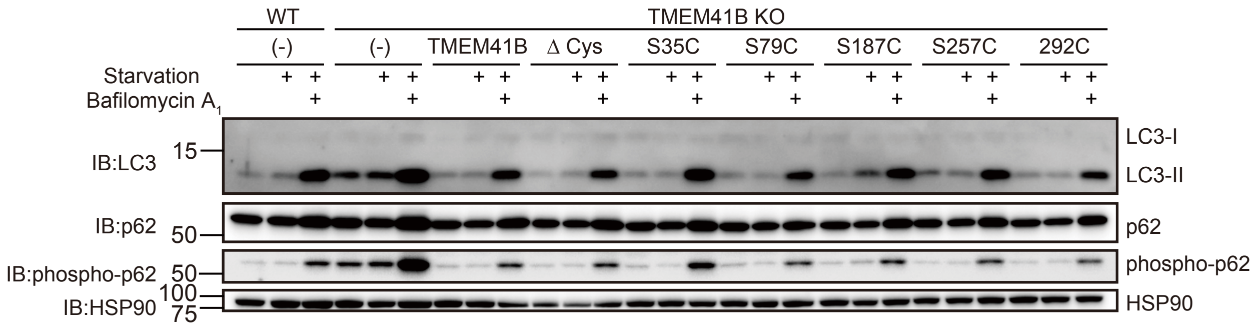

**Figure S5.** The single cysteine TMEM41B mutants retained autophagic function. TMEM41B-KO HeLa cells stably expressing cysteine-less TMEM41B (ΔCys) or the single cysteine mutants were starved with or without bafilomycin A1 treatment. Cell lysates were subjected to immunoblotting using anti-p62, anti-phospho-p62, and anti-LC3 antibodies.

Table S1. DedA superfamily proteins analyzed in Fig. 2.

| Number (*1) | Sequence name | Family name | Taxonomy | Note |
|---|---|---|---|---|
| 1 | Candidatus_syntrophicum_VTT_domain_containing_protein_WP_147662241 | VMP1 | Archaea | |
| 2 | Candidatus_syntrophicum_SNARE_associated_Golgi_protein_QEE16945 | TMEM41 | Archaea | |
| 3 | Candidatus_syntrophicum_hypothetical_protein_WP_147664438 | TMEM41 | Archaea | |
| 4 | Arabidopsis_thaliana_SNARE_associated_Golgi_protein_family_NP_172707 | TMEM41 | Eukarya | |
| 5 | Arabidopsis_thaliana_small_multi_drug_export_protein_NP_178363 | PF06695 | Eukarya | |
| 6 | Arabidopsis_thaliana_SNARE_associated_Golgi_protein_family_NP_192937 | TMEM41 | Eukarya | |
| 7 | Helicobacter_pylori_26695_membrane_protein_NP_207366 | DedA | Bacteria | |
| 8 | Helicobacter_pylori_26695_membrane_protein_NP_207953 | DedA | Bacteria | |
| 9 | Neisseria_meningitidis_MC58_hypothetical_protein_NMB0534_NP_273579 | DedA | Bacteria | |
| 10 | Neisseria_meningitidis_MC58_dedA_protein_NP_274086 | DedA | Bacteria | |
| 11 | Neisseria_meningitidis_MC58_dedA_protein_NP_274693 | DedA | Bacteria | |
| 12 | Bacillus_subtilis_subsp_subtilis_str_168_putative_membrane_phosphatase_NP_388110 | DedA | Bacteria | |
| 13 | Bacillus_subtilis_subsp_subtilis_str_168_putative_integral_membrane_protein_NP_388929 | TMEM41 | Bacteria | |
| 14 | Bacillus_subtilis_subsp_subtilis_str_168_putative_integral_membrane_protein_NP_389226 | DedA | Bacteria | |
| 15 | Bacillus_subtilis_subsp_subtilis_str_168_putative_integral_inner_membrane_protein_phosphatase_or_phosphate_isomerase_NP_389701 | DedA | Bacteria | |
| 16 | Bacillus_subtilis_subsp_subtilis_str_168_conserved_membrane_protein_of_unknown_function_NP_390449 | TMEM41 | Bacteria | |
| 17 | Bacillus_subtilis_subsp_subtilis_str_168_putative_osomosensing_transporter_NP_390775 | TMEM41 | Bacteria | |
| 18 | Escherichia_coli_str_K_12_substr_MG1655_DedA_family_protein_YabI_NP_414607 | DedA | Bacteria | EcYabI |
| 19 | Escherichia_coli_str_K_12_substr_MG1655_DedA_family_protein_YdjZ_NP_416266 | TMEM41 | Bacteria | EcYdjZ |
| 20 | Escherichia_coli_str_K_12_substr_MG1655_DedA_family_protein_DedA_NP_416820 | DedA | Bacteria | EcDedA |
| 21 | Escherichia_coli_str_K_12_substr_MG1655_DedA_family_protein_YqaA_NP_417174 | DedA | Bacteria | EcYqaA |
| 22 | Escherichia_coli_str_K_12_substr_MG1655_DedA_family_protein_YghB_NP_417482 | DedA | Bacteria | EcYghB |
| 23 | Escherichia_coli_str_K_12_substr_MG1655_DedA_family_protein_YqjA_NP_417566 | DedA | Bacteria | EcYqjA |
| 24 | Caenorhabditis_elegans_Transmembrane_protein_41_homolog_NP_495985 | TMEM41 | Eukarya | |
| 25 | Homo_sapiens_transmembrane_protein_41A_precursor_NP_542383 | TMEM41 | Eukarya | HsTMEM41A |
| 26 | Arabidopsis_thaliana_vacuole_membrane_like_protein_NP_563735 | VMP1 | Eukarya | |
| 27 | Arabidopsis_thaliana_SNARE_associated_Golgi_protein_family_NP_564182 | TMEM41 | Eukarya | |
| 28 | Arabidopsis_thaliana_SNARE_associated_Golgi_protein_family_NP_565283 | TMEM41 | Eukarya | |
| 29 | Arabidopsis_thaliana_SNARE_associated_Golgi_protein_family_NP_565028 | TMEM41 | Eukarya | |
| 30 | Arabidopsis_thaliana_SNARE_associated_Golgi_protein_family_NP_567450 | VMP1 | Eukarya | |
| 31 | Arabidopsis_thaliana_SNARE_associated_Golgi_protein_family_NP_567541 | TMEM41 | Eukarya | |
| 32 | Drosophila_melanogaster_stasimon_isoform_A_NP_573225 | TMEM41 | Eukarya | |
| 33 | Homo_sapiens_vacuole_membrane_protein_1_isoform_1_NP_112200 | VMP1 | Eukarya | HsVMP1 |
| 34 | Drosophila_melanogaster_uncharacterized_protein_Dmel_CG11367_NP_649412 | TMEM41 | Eukarya | |
| 35 | Arabidopsis_thaliana_SNARE_associated_Golgi_protein_family_NP_175116 | TMEM41 | Eukarya | |
| 36 | Drosophila_melanogaster_transport_and_golgi_organization_5_isoform_A_NP_727444 | VMP1 | Eukarya | |
| 37 | Drosophila_melanogaster_uncharacterized_protein_Dmel_CG32087_NP_729740 | VMP1 | Eukarya | |
| 38 | Arabidopsis_thaliana_SNARE_associated_Golgi_protein_family_NP_192696 | TMEM41 | Eukarya | |
| 39 | Arabidopsis_thaliana_SNARE_associated_Golgi_protein_family_NP_194016 | TMEM41 | Eukarya | |
| 40 | Arabidopsis_thaliana_SNARE_associated_Golgi_protein_family_NP_197408 | TMEM41 | Eukarya | |
| 41 | Caenorhabditis_elegans_Ectopic_P_granules_protein_3_NP_499688 | VMP1 | Eukarya | |
| 42 | Arabidopsis_thaliana_SNARE_associated_Golgi_protein_family_NP_171825 | TMEM41 | Eukarya | |
| 43 | Thermus_thermophilus_HB8_DedA_family_protein_YP_143783 | DedA | Bacteria | |
| 44 | Thermus_thermophilus_HB8_small_multidrug_export_protein_YP_143820 | PF06695 | Bacteria | |
| 45 | Homo_sapiens_transmembrane_protein_41B_isoform_1_NP_055827 | TMEM41 | Eukarya | HsTMEM41B |
| 46 | Entamoeba_histolytica_HM_1_IMSS_hypothetical_protein_EHI_169620_XP_650378 | other | Eukarya | |
| 47 | Entamoeba_histolytica_HM_1_IMSS_hypothetical_protein_EHI_049010_XP_651073 | TMEM41 | Eukarya | |
| 48 | Entamoeba_histolytica_HM_1_IMSS_hypothetical_protein_conserved_XP_654786 | TMEM41 | Eukarya | |
| 49 | Entamoeba_histolytica_HM_1_IMSS_hypothetical_protein_EHI_027530_XP_654900 | VMP1 | Eukarya | |
| 50 | Escherichia_coli_str_K_12_substr_MG1655_DedA_family_protein_YdjX_NP_416264 | TMEM41 | Bacteria | EcYdjX |
| 51 | Escherichia_coli_str_K_12_substr_MG1655_DedA_family_protein_YohD_NP_416640 | DedA | Bacteria | EcYohD |
| 52 | Caenorhabditis_elegans_Uncharacterized_protein_CELE_T07F104_NP_001041162 | TMEM41 | Eukarya | |
| 53 | Caenorhabditis_elegans_Uncharacterized_protein_CELE_Y71A12C2_NP_493447 | TMEM41 | Eukarya | |
| 54 | Plasmodium_falciparum_3D7_SNARE_associated_Golgi_protein_putative_XP_001350367 | TMEM41 | Eukarya | |
| 55 | Plasmodium_falciparum_3D7_conserved_Plasmodium_membrane_protein_unknown_function_XP_001348888 | VMP1 | Eukarya | |
| 56 | Entamoeba_histolytica_HM_1_IMSS_uncharacterized_protein_EHI_108350_XP_655504 | TMEM41 | Eukarya | |
| 57 | Yersinia_pestis_CO92_DedA_family_membrane_protein_YP_002345593 | DedA | Bacteria | |
| 58 | Yersinia_pestis_CO92_DedA_family_membrane_protein_YP_002345646 | DedA | Bacteria | |
| 59 | Yersinia_pestis_CO92_DedA_family_membrane_protein_YP_002345746 | DedA | Bacteria | |
| 60 | Yersinia_pestis_CO92_hypothetical_protein_YPO2767_YP_002347714 | DedA | Bacteria | |
| 61 | Yersinia_pestis_CO92_hypothetical_protein_YPO3302_YP_002348198 | DedA | Bacteria | |
| 62 | Yersinia_pestis_CO92_hypothetical_protein_YPO3783_YP_002348660 | DedA | Bacteria | |
| 63 | Homo_sapiens_transmembrane_protein_64_isoform_1_NP_001008495 | TMEM41 | Eukarya | HsTMEM64 |
| 64 | Homo_sapiens_transmembrane_protein_64_isoform_2_NP_001139745 | TMEM41 | Eukarya | |
| 65 | Arabidopsis_thaliana_SNARE_associated_Golgi_protein_family_NP_001154265 | TMEM41 | Eukarya | |
| 66 | Arabidopsis_thaliana_transmembrane_protein_NP_193051 | DedA | Eukarya | |
| 67 | Plasmodium_falciparum_3D7_SNARE_associated_Golgi_protein_putative_XP_966099 | TMEM41 | Eukarya | |
| 68 | Arabidopsis_thaliana_SNARE_associated_Golgi_protein_family_NP_001185376 | TMEM41 | Eukarya | |
| 69 | Saccharomyces_cerevisiae_S288C_Tvp38p_NP_013014 | TMEM41 | Eukarya | ScTvp38 |
| 70 | Schizosaccharomyces_pombe_putative_SNARE_associated_protein_NP_595882 | TMEM41 | Eukarya | |
| 71 | Solanum_lycopersicum_uncharacterized_protein_LOC101257596_XP_004229441 | TMEM41 | Eukarya | |
| 72 | Solanum_lycopersicum_uncharacterized_protein_LOC101246475_XP_004230383 | TMEM41 | Eukarya | |
| 73 | Solanum_lycopersicum_uncharacterized_protein_LOC101255587_XP_004238763 | PF06695 | Eukarya | |
| 74 | Solanum_lycopersicum_uncharacterized_membrane_protein_At4g09580_XP_004241822 | TMEM41 | Eukarya | |
| 75 | Solanum_lycopersicum_uncharacterized_protein_LOC101245369_XP_004241892 | TMEM41 | Eukarya | |
| 76 | Solanum_lycopersicum_transmembrane_protein_64_XP_004245695 | TMEM41 | Eukarya | |
| 77 | Solanum_lycopersicum_uncharacterized_membrane_protein_At4g09580_XP_004245963 | TMEM41 | Eukarya | |
| 78 | Solanum_lycopersicum_uncharacterized_protein_LOC101255624_XP_004247084 | DedA | Eukarya | |
| 79 | Solanum_lycopersicum_vacuole_membrane_protein_KMS1_XP_004247368 | VMP1 | Eukarya | |
| 80 | Solanum_lycopersicum_uncharacterized_membrane_protein_At4g09580_XP_004250397 | TMEM41 | Eukarya | |
| 81 | Solanum_lycopersicum_uncharacterized_protein_LOC101260690_XP_004250700 | TMEM41 | Eukarya | |
| 82 | Sulfolobaceae_MULTISPECIES__hypothetical_protein_WP_009992141 | DedA | Archaea | |
| 83 | Archaeoglobus_fulgidus_COG2426_family_protein_WP_010879192 | PF06695 | Archaea | |
| 84 | Methanosarcina_MULTISPECIES__small_multi_drug_export_protein_WP_011033760 | PF06695 | Archaea | |
| 85 | Streptomyces_scabiei_DedA_family_protein_WP_012999571 | DedA | Bacteria | |
| 86 | Streptomyces_scabiei_membrane_protein_WP_013001687 | DedA | Bacteria | |
| 87 | Streptomyces_scabiei_DedA_family_protein_WP_013004004 | DedA | Bacteria | |
| 88 | Streptomyces_scabiei_membrane_protein_WP_013004015 | DedA | Bacteria | |
| 89 | Streptomyces_scabiei_TVP38_TMEM64_family_protein_WP_013005183 | TMEM41 | Bacteria | |
| 90 | Streptomyces_scabiei_DedA_family_protein_WP_013005189 | DedA | Bacteria | |
| 91 | Fischerella_muscicola_DedA_family_protein_WP_016865728 | DedA | Bacteria | |
| 92 | Fischerella_muscicola_DedA_family_protein_WP_016867571 | DedA | Bacteria | |
| 93 | Fischerella_muscicola_TVP38_TMEM64_family_protein_WP_016870429 | TMEM41 | Bacteria | |
| 94 | Fischerella_muscicola_TVP38_TMEM64_family_protein_WP_016870430 | TMEM41 | Bacteria | |
| 95 | Ustilago_maydis_521_hypothetical_protein_UMAG_00272_XP_011386189 | TMEM41 | Eukarya | |
| 96 | Ustilago_maydis_521_hypothetical_protein_UMAG_00958_XP_011386954 | TMEM41 | Eukarya | |
| 97 | Ustilago_maydis_521_hypothetical_protein_UMAG_01159_XP_011387102 | TMEM41 | Eukarya | |

| | | | |
|---|---|---|---|
| 98 | Ustilago_maydis_521_hypothetical_protein_UMAG_05456_XP_011391801 | TMEM41 | Eukarya |
| 99 | Ustilago_maydis_521_hypothetical_protein_UMAG_06420_XP_011392703 | TMEM41 | Eukarya |
| 100 | Methanosarcina_mazei_DedA_family_protein_WP_048046344 | DedA | Archaea |
| 101 | Pyrococcus_horikoshii_COG2426_family_protein_WP_048053449 | PF06695 | Archaea |
| 102 | Tenacibaculum_dicentrarchi_short_chain_dehydrogenase_WP_058885722 | DedA | Bacteria |
| 103 | Homo_sapiens_transmembrane_protein_41A_isoform_X1_XP_016862926 | TMEM41 | Eukarya |
| 104 | Solanum_lycopersicum_vacuole_membrane_protein_KMS1_like_NP_001315551 | VMP1 | Eukarya |
| 105 | Arabidopsis_thaliana_SNARE_associated_Golgi_protein_family_NP_001331805 | TMEM41 | Eukarya |
| 106 | Clostridium_sp_Pfam_Sm_multidrug_ex_R7M7P8 | PF06695 | Bacteria |
| 107 | Crocosphaera_subtropica_Pfam_SNARE_VTT_B1WQI7 | TMEM41 | Bacteria |
| 108 | Clostridium_sp_Uncharacterized_protein_R6XST9 | PF06695 | Bacteria |
| 109 | Eubacterium_sp_Uncharacterized_protein_R6QNG8 | PF06695 | Bacteria |
| 110 | Faecalibacterium_sp_Small_multi_drug_export_protein_R5FIZ3 | PF06695 | Bacteria |
| 111 | Thalassiosira_oceanica_Uncharacterized_protein_Fragment_K0TKX5 | PF06695 | Eukarya |
| 112 | Clostridium_sp_Putative_small_multi_drug_export_protein_R5K8E9 | PF06695 | Bacteria |
| 113 | Xenopus_tropicalis_Cytoskeleton_associated_protein_2_like_F6U3L8 | other | Eukarya |
| 114 | Herpetosiphon_aurantiacus_SNARE_associated_Golgi_protein_A9B4D4 | VMP1 | Bacteria |
| 115 | Alkaliphilus_metalliredigens_TVP38_TMEM64_family_membrane_protein_A6TJC3 | TMEM41 | Bacteria |
| 116 | Methylorubrum_extorquens_TVP38_TMEM64_family_membrane_protein_C5ARL8 | TMEM41 | Bacteria |
| 117 | Aeromonas_hydrophila_DedA_family_protein_A0KPF7 | DedA | Bacteria |

(*1) The numbers correspond to those in Fig. 2B

Table S2. Transmembrane proteins containing 2 loops found in PDBTM (*1)

| PDB ID | Protein | facing | Note |
|---|---|---|---|
| 1fqy | AQUAPORIN-1 | Yes | turning with loop structure |
| 1fx8 | GLYCEROL FACILITATOR (GLPF) WITH SUBSTRATE GLYCEROL | Yes | turning with loop structure |
| 1h6i | AQUAPORIN 1 | Yes | turning with loop structure |
| 1ih5 | AQUAPORIN-1 | Yes | turning with loop structure |
| 1j4n | AQP1 WATER CHANNEL | Yes | turning with loop structure |
| 1kpk | CLC CHLORIDE CHANNEL | Yes | Passing each other, Forming a complex with the RE inside |
| 1kpl | CLC CHLORIDE CHANNEL | Yes | Passing each other, Forming a complex with the RE inside |
| 1lda | GLYCEROL FACILITATOR (GLPF) | Yes | turning with loop structure |
| 1ldf | GLYCEROL FACILITATOR (GLPF) MUTATION W48F, F200T | Yes | turning with loop structure |
| 1ldi | GLYCEROL FACILITATOR (GLPF) WITHOUT SUBSTRATE GLYCEROL | Yes | turning with loop structure |
| 1ntk | MITOCHONDRIAL CYTOCHROME BC1 IN COMPLEX WITH ANTIMYCIN A1 | No | |
| 1ots | CLC CHLORIDE CHANNEL AND FAB COMPLEX | Yes | Passing each other, Forming a complex with the RE inside |
| 1ott | CLC CHLORIDE CHANNEL E148A MUTANT AND FAB COMPLEX | Yes | Passing each other, Forming a complex with the RE inside |
| 1otu | CLC CHLORIDE CHANNEL E148Q MUTANT AND FAB COMPLEX | Yes | Passing each other, Forming a complex with the RE inside |
| 1rc2 | AQUAPORIN Z | Yes | turning with loop structure |
| 1s6e | PORIN 6 | Yes | turning with loop structure |
| 1sor | AQUAPORIN-0 CLOSED WATER PORE | Yes | turning with loop structure |
| 1ufd | NODULIN 26 | Yes | turning with loop structure |
| 1xfh | GLUTAMATE TRANSPORTER | Yes | |
| 1ymg | AQUAPORIN O | Yes | turning with loop structure |
| 1z98 | AQUAPORIN SOPIP2 | Yes | turning with loop structure |
| 2abm | AQUAPORIN Z | Yes | turning with loop structure |
| 2b5f | AQUAPORIN SOPIP2;1 IN AN OPEN CONFORMATION | Yes | turning with loop structure |
| 2b6o | LENS AQUAPORIN-0 IN A CLOSED PORE STATE | Yes | turning with loop structure |
| 2b6p | LENS AQUAPORIN-0 (AQP0) (LENS MIP) IN AN OPEN PORE STATE | Yes | turning with loop structure |
| 2c32 | EYE LENS AQUAPORIN-0 | Yes | turning with loop structure |
| 2d57 | AQUAPORIN-4 (AQP4M23) | Yes | turning with loop structure |
| 2evu | AQUAPORIN AQPM | Yes | turning with loop structure |
| 2exw | ECCLC-FAB COMPLEX IN THE ABSENCE OF BOUND IONS | Yes | Passing each other, Forming a complex with the RE inside |
| 2exy | E148Q MUTANT OF ECCLC, FAB COMPLEXED IN ABSENCE OF BOUND IONS | Yes | Passing each other, Forming a complex with the RE inside |
| 2ez0 | S107A/E148Q/Y445A MUTANT OF ECCLC, IN COMPLEX WITH A FAB FRAGMENT | Yes | Passing each other, Forming a complex with the RE inside |
| 2f2b | AQUAPORIN AQPM | Yes | turning with loop structure |
| 2fec | E203Q MUTANT OF THE CL-/H+ EXCHANGER CLC- EC1 | Yes | Passing each other, Forming a complex with the RE inside |
| 2fed | E203Q MUTANT OF THE CL-/H+ EXCHANGER CLC- EC1 | Yes | Passing each other, Forming a complex with the RE inside |
| 2fee | CL-/H+ EXCHANGER CLC-EC1 | Yes | Passing each other, Forming a complex with the RE inside |
| 2h2p | CLC-EC1 IN COMPLEX WITH FAB FRAGMENT IN SECN- | Yes | Passing each other, Forming a complex with the RE inside |
| 2h2s | E148A MUTANT OF CLC-EC1 IN SECN- | Yes | Passing each other, Forming a complex with the RE inside |
| 2hlf | CLC CHLORIDE CHANNEL Y445E MUTANT AND FAB COMPLEX | Yes | Passing each other, Forming a complex with the RE inside |
| 2ht2 | CLC CHLORIDE CHANNEL Y445H MUTANT AND FAB COMPLEX | Yes | Passing each other, Forming a complex with the RE inside |
| 2ht3 | CLC CHLORIDE CHANNEL Y445L MUTANT AND FAB COMPLEX | Yes | Passing each other, Forming a complex with the RE inside |
| 2ht4 | CLC CHLORIDE CHANNEL Y445W MUTANT AND FAB COMPLEX | Yes | Passing each other, Forming a complex with the RE inside |
| 2htk | CLC CHLORIDE CHANNEL Y445A MUTANT AND FAB COMPLEX | Yes | Passing each other, Forming a complex with the RE inside |
| 2htl | CLC CHLORIDE CHANNEL Y445F MUTANT AND FAB COMPLEX | Yes | Passing each other, Forming a complex with the RE inside |
| 2nwl | GLTPH IN COMPLEX WITH L-ASP | Yes | |
| 2nww | GLTPH IN COMPLEX WITH TBOA | Yes | |
| 2nwx | GLTPH IN COMPLEX WITH L-ASPARTATE AND SODIUM IONS | Yes | |
| 2o9d | AQPZ MUTANT T183C | Yes | turning with loop structure |
| 2o9e | AQPZ MUTANT T183C COMPLEXED WITH MERCURY | Yes | turning with loop structure |
| 2o9f | AQPZ MUTANT L170C | Yes | turning with loop structure |
| 2o9g | AQPZ MUTANT L170C COMPLEXED WITH MERCURY | Yes | turning with loop structure |
| 2r9h | Q207C MUTANT OF CLC-EC1 IN COMPLEX WITH FAB | Yes | Passing each other, Forming a complex with the RE inside |
| 2w1p | P.PASTORIS AQUAPORIN | Yes | turning with loop structure |
| 2w2e | P.PASTORIS AQUAPORIN | Yes | turning with loop structure |
| 2zz9 | AQUAPORIN-4 S180D MUTANT | Yes | turning with loop structure |
| 3c02 | AQUAGLYCEROPORIN | Yes | turning with loop structure |
| 3cll | AQUAPORIN SOPIP2;1 S115E MUTANT | Yes | turning with loop structure |
| 3cn5 | AQUAPORIN SOPIP2;1 S115E, S274E MUTANT | Yes | turning with loop structure |
| 3cn6 | AQUAPORIN SOPIP2 | Yes | turning with loop structure |
| 3d9s | AQUAPORIN 5 (AQP5) | Yes | turning with loop structure |
| 3det | E418A, Y445A DOUBLY UNGATED MUTANT OF E.COLI CLC_EC1 | Yes | Passing each other, Forming a complex with the RE inside |
| 3ejy | E203H MUTANT OF E.COLI CL-/H+ ANTIPORTER, CLC- EC1 | Yes | Passing each other, Forming a complex with the RE inside |
| 3ejz | E203V MUTANT E.COLI CL-/H+ EXCHANGER, CLC-EC1 | Yes | Passing each other, Forming a complex with the RE inside |
| 3gd8 | AQUAPORIN 4 | Yes | turning with loop structure |
| 3j41 | AQUAPORIN-0/CALMODULIN COMPLEX | Yes | turning with loop structure |
| 3jav | IP3R1 CHANNEL IN THE APO-STATE | No | |
| 3k3g | UREA TRANSPORTER | Yes | turning with loop structure |
| 3llq | AQUAPORIN | Yes | turning with loop structure |
| 3m6e | UREA TRANSPORTER | Yes | turning with loop structure |
| 3m9i | LENS AQUAPORIN-0 (AQP0) | Yes | turning with loop structure |
| 3nd0 | SLOW CYANOBACTERIAL CL-/H+ ANTIPORTER | Yes | Passing each other, Forming a complex with the RE inside |
| 3ne2 | AQUAPORIN | Yes | turning with loop structure |
| 3nk5 | AQPZ MUTANT F43W | Yes | turning with loop structure |
| 3nka | AQPZ H174G,T183F | Yes | turning with loop structure |
| 3nkc | AQPZ F43W,H174G,T183F | Yes | turning with loop structure |
| 3nmo | CLC-EC1 CL-/H+ TRANSPORTER | Yes | Passing each other, Forming a complex with the RE inside |
| 3org | EUKARYOTIC CLC TRANSPORTER | Yes | Passing each other, Forming a complex with the RE inside |
| 3q17 | SLOW CLC CL-/H+ ANTIPORTER | Yes | Passing each other, Forming a complex with the RE inside |
| 3tdo | HSC | Yes | turning with loop structure |
| 3tdp | HSC | Yes | turning with loop structure |
| 3tdr | HSC | Yes | turning with loop structure |
| 3tds | HSC F194I | Yes | turning with loop structure |
| 3tdx | HSC L82V | Yes | turning with loop structure |
| 3te0 | HSC K148E | Yes | turning with loop structure |
| 3te1 | HSC T84A | Yes | turning with loop structure |
| 3te2 | HSC K16S | Yes | turning with loop structure |
| 3ukm | TWO PORE DOMAIN POTASSIUM ION CHANNEL K2P1 (TWIK-1) | Yes | Parallel, Forming a complex with the RE inside |
| 3um7 | TWO PORE DOMAIN K+ ION CHANNEL TRAAK (K2P4.1) | Yes | Parallel, Forming a complex with the RE inside, turning with loop structure |
| 3v8g | ASYMMETRIC TRIMER OF A GLUTAMATE TRANSPORTER | Yes | |
| 3zoj | AQUAPORIN AQY1 | Yes | turning with loop structure |
| 4bw5 | TWO PORE DOMAIN POTASSIUM ION CHANNEL TREK2 (K2P10.1) | Yes | Passing each other, Forming a complex with the RE inside |
| 4csk | AQUAPORIN | Yes | turning with loop structure |
| 4ene | N- AND C-TERMINAL TRIMMED CLC-EC1 CL-/H+ ANTIPORTER AND FAB COMPLEX | Yes | Passing each other, Forming a complex with the RE inside |
| 4ezc | UREA TRANSPORTER | Yes | turning with loop structure |
| 4ezd | UREA TRANSPORTER BOUND TO SELENOUREA | Yes | turning with loop structure |
| 4fc4 | FNT FAMILY ION CHANNEL | Yes | turning with loop structure |
| 4fg6 | ECCLC E148A MUTANT | Yes | Passing each other, Forming a complex with the RE inside |
| 4ftp | E202Y MUTANT OF THE CL-/H+ ANTIPORTER CLC-EC1 | Yes | Passing each other, Forming a complex with the RE inside |
| 4g1u | HEME TRANSPORTER HMUUV | Yes | turning with loop structure |
| 4i9w | TWO PORE DOMAIN K+ CHANNEL TRAAK (K2P4.1) - FAB COMPLEX | Yes | Parallel, Forming a complex with the RE inside, turning with loop structure |
| 4ia4 | AQUAPORIN SOPIP2 | Yes | turning with loop structure |
| 4izm | GLTPH L66C-S300C MUTANT CROSSLINKED WITH DIVALENT MERCURY | Yes | |
| 4j7c | POTASSIUM TRANSPORTER | Yes | Parallel, turning with loop structure |
| 4jc6 | PLANT AQUAPORIN SOPIP2;1 | Yes | turning with loop structure |
| 4kjp | CLC-EC1 DELTANC CONSTRUCT IN THE ABSENCE OF HALIDE | Yes | Passing each other, Forming a complex with the RE inside |
| 4kjq | CLC-EC1 DELTANC CONSTRUCT IN 100MM FLUORIDE | Yes | Passing each other, Forming a complex with the RE inside |
| 4kjw | CLC-EC1 DELTANC CONSTRUCT IN 100MM FLUORIDE AND 20MM BROMIDE | Yes | Passing each other, Forming a complex with the RE inside |
| 4kk5 | CLC-EC1 DELTANC CONSTRUCT IN 20MM FLUORIDE AND 20MM BROMIDE | Yes | Passing each other, Forming a complex with the RE inside |
| 4kk6 | CLC-EC1 DELTANC CONSTRUCT IN 20MM BROMIDE | Yes | Passing each other, Forming a complex with the RE inside |
| 4kk8 | E148Q MUTANT OF CLC-EC1 DELTANC CONSTRUCT IN 100MM FLUORIDE | Yes | Passing each other, Forming a complex with the RE inside |

| | | | |
|---|---|---|---|
| 4kk9 | E148A MUTANT OF CLC-EC1 DELTANC CONSTRUCT IN 100MM FLUORIDE AND 2MM | Yes | Passing each other, Forming a complex with the RE inside |
| 4kka | E148A MUTANT OF CLC-EC1 DELTANC CONSTRUCT IN 100MM FLUORIDE AND 20MM BROMIDE | Yes | Passing each other, Forming a complex with the RE inside |
| 4kkb | E148A MUTANT OF CLC-EC1 DELTANC CONSTRUCT IN 20MM FLUORIDE AND 20MM BROMIDE | Yes | Passing each other, Forming a complex with the RE inside |
| 4kkc | E148A MUTANT OF CLC-EC1 DELTANC CONSTRUCT IN 20MM BROMIDE | Yes | Passing each other, Forming a complex with the RE inside |
| 4kkl | E148A MUTANT OF CLC-EC1 DELTA NC CONSTRUCT IN 100MM FLUORIDE | Yes | Passing each other, Forming a complex with the RE inside |
| 4ky0 | SUBSTRATE-FREE GLUTAMATE TRANSPORTER | Yes | |
| 4lou | E148Q MUTANT OF CLC-EC1 DELTANC CONSTRUCT IN THE ABSENCE OF HALIDE | Yes | Passing each other, Forming a complex with the RE inside |
| 4mqx | CLC-EC1 FAB COMPLEX CYSLESS A399C-A432C MUTANT | Yes | Passing each other, Forming a complex with the RE inside |
| 4nef | AQUAPORIN 2 | Yes | turning with loop structure |
| 4oj2 | AQUAPORIN | Yes | turning with loop structure |
| 4oye | GLTPH R397A IN APO | Yes | |
| 4rue | K2P4.1 (TRAAK) POTASSIUM CHANNEL, G124I MUTANT | Yes | Parallel, Forming a complex with the RE inside, turning with loop structure |
| 4ruf | K2P4.1 (TRAAAK) POTASSIUM CHANNEL, W262S MUTAN | Yes | Parallel, Forming a complex with the RE inside, turning with loop structure |
| 4twk | POTASSIUM ION CHANNEL TREK1 (K2P2.1) | Yes | Parallel, Forming a complex with the RE inside, turning with loop structure |
| 4wfe | TRAAK K+ CHANNEL IN A K+ BOUND CONDUCTIVE CONFORMATION | Yes | Parallel, Forming a complex with the RE inside, turning with loop structure |
| 4wff | TRAAK K+ CHANNEL IN A K+ BOUND NONCONDUCTIVE CONFORMATION | Yes | Parallel, Forming a complex with the RE inside, turning with loop structure |
| 4wfg | TRAAK K+ CHANNEL IN A TL+ BOUND CONDUCTIVE CONFORMATION | Yes | Parallel, Forming a complex with the RE inside, turning with loop structure |
| 4wfh | TRAAK K+ CHANNEL IN A TL+ BOUND NONCONDUCTIVE CONFORMATION | Yes | Parallel, Forming a complex with the RE inside, turning with loop structure |
| 4xdj | POTASSIUM ION CHANNEL TREK2 (K2P10.1) IN AN ALTERNATE CONFORMATION (FORM 2) | Yes | Parallel, Forming a complex with the RE inside, turning with loop structure |
| 4xdl | POTASSIUM ION CHANNEL TREK2 (K2P10.1) IN COMPLEX WITH A BROMINATED FLUOXETINE DERIVATIVE. | Yes | Parallel, Forming a complex with the RE inside, turning with loop structure |
| 5a1s | SODIUM-DEPENDENT CITRATE SYMPORTER SECITS | Yes | |
| 5a6u | RIBOSOME-BOUND SEC61 PROTEIN-CONDUCTING CHANNEL IN THE 'NON-INSERTING' STATE | No | |
| 5bn2 | ROOM TEMPERATURE STRUCTURE OF AQUAPORIN | Yes | turning with loop structure |
| 5c5x | S156E MUTANT OF HUMAN AQUAPORIN 5 | Yes | turning with loop structure |
| 5cfy | GLTPH R397A IN COMPLEX WITH NA+ AND L-ASP | Yes | |
| 5dwy | SUBSTRATE-FREE GLUTAMATE TRANSPORTER HOMOLOGUE GLTTK | Yes | |
| 5e1j | VOLTAGE-GATED TWO-PORE CHANNEL TPC1 | Yes | |
| 5e9s | SUBSTRATE-BOUND GLUTAMATE TRANSPORTER HOMOLOGUE GLTTK | Yes | |
| 5fvn | OMPE36 PORIN | β strand | |
| 5i32 | AMMONIA PERMEABLE AQUAPORIN ATTIP2;1 | Yes | turning with loop structure |
| 5iws | TRANSPORTER MALT, THE EIIC DOMAIN | Yes | |
| 5l2a | CNTNW N149S,F366A IN AN OUTWARD-FACING STATE | No | |
| 5l2b | CNTNW N149S, E332A IN AN OUTWARD-FACING STATE | No | |
| 5mg3 | BACTERIAL HOLO-TRANSLOCON | Yes | |
| 5oon | UNDECAPRENYL-PYROPHOSPHATE PHOSPHATASE, BACA | Yes | |
| 5tqq | CLC-K CHLORIDE CHANNEL, MAIN (CLASS 1) CONFORMATION | Yes | |
| 5tua | NA+-SELECTIVE MUTANT OF TWO-PORE CHANNEL | Yes | Parallel, Forming a complex with the RE inside, turning with loop structure |
| 5u9w | CNTNW N149L IN THE INTERMEDIATE 3 STATE | No | |
| 5x9r | ELEVATOR-LIKE MECHANISM OF THE SODIUM/CITRATE SYMPORTER CITS | Yes | |
| 5xar | SODIUM/CITRATE SYMPORTER CITS | Yes | |
| 5xjy | ABCA1 | No | |
| 5zdh | ETEC PILOTIN-SECRETIN ASPS-GSPD COMPLEX | β strand | |
| 5zx5 | TRPM7 WITH EDTA | No | |
| 6ad7 | E148D MUTANT CLC-EC1 IN 20 MM BROMIDE | Yes | Passing each other, Forming a complex with the RE inside |
| 6ad8 | E148D MUTANT CLC-EC1 IN 50 MM BROMIDE | Yes | Passing each other, Forming a complex with the RE inside |
| 6ada | E148D MUTANT CLC-EC1 IN 200MM BROMIDE | Yes | Passing each other, Forming a complex with the RE inside |
| 6adb | E148N MUTANT CLC-EC1 IN 20MM BROMIDE | Yes | Passing each other, Forming a complex with the RE inside |
| 6adc | E148A MUTANT CLC-EC1 IN THE PRESENCE OF 50MM BROMOACETATE | Yes | Passing each other, Forming a complex with the RE inside |
| 6bat | GLTPH IN COMPLEX WITH L-ASPARTATE | Yes | |
| 6bau | GLTPH R397C IN COMPLEX WITH L-CYSTEINE | Yes | |
| 6bav | GLTPH R397C IN COMPLEX WITH S-BENZYL-L-CYSTEINE | Yes | |
| 6bmi | GLTPH R397C IN COMPLEX WITH L-SERINE | Yes | |
| 6bqr | TRPM4 ION CHANNEL IN LIPID NANODISCS IN A CALCIUM-FREE STATE | Yes | Parallel, Forming a complex with the RE inside, turning with loop structure |
| 6c96 | TPC1 CHANNEL | Yes | Parallel, Forming a complex with the RE inside, turning with loop structure |
| 6c9a | MOUSE TPC1 CHANNEL IN THE PTDINS(3,5)P2-BOUND STATE | Yes | Parallel, Forming a complex with the RE inside, turning with loop structure |
| 6cb2 | UPPP | Yes | |
| 6cq6 | K2P2.1(TREK-1) APO | Yes | Parallel, Forming a complex with the RE inside, turning with loop structure |
| 6cq8 | K2P2.1(TREK-1):ML335 COMPLEX | Yes | Parallel, Forming a complex with the RE inside, turning with loop structure |
| 6cq9 | K2P2.1(TREK-1):ML402 COMPLEX | Yes | Parallel, Forming a complex with the RE inside, turning with loop structure |
| 6cx0 | ATTPC1 D376A | Yes | Parallel, Forming a complex with the RE inside, turning with loop structure |
| 6drj | TRPM2 ION CHANNEL RECEPTOR BY SINGLE PARTICLE ELECTRON CRYO-MICROSCOPY, ADPR/CA2+ BOUND STATE | Yes | Parallel, Forming a complex with the RE inside, turning with loop structure |
| 6e1k | ATTPC1(DDE) RECONSTITUTED IN SAPOSIN A WITH CAT06 FAB | Yes | Parallel, Forming a complex with the RE inside, turning with loop structure |
| 6e1m | ATTPC1(DDE) RECONSTITUTED IN SAPOSIN A | Yes | Parallel, Forming a complex with the RE inside, turning with loop structure |
| 6e1n | ATTPC1(DDE) IN STATE 1 | Yes | Parallel, Forming a complex with the RE inside, turning with loop structure |
| 6e1p | ATTPC1(DDE) IN STATE 2 | Yes | Parallel, Forming a complex with the RE inside, turning with loop structure |
| 6f7h | AQP10 | Yes | turning with loop structure |
| 6k5a | E148D/R147A/F317A MUTANT IN PRESENCE OF 200 MM NABR | Yes | Passing each other, Forming a complex with the RE inside |
| 6k5d | E148N MUTANT CLC-EC1 IN PRESENCE OF 200 MM NABR | Yes | Passing each other, Forming a complex with the RE inside |
| 6k5f | CLC-EC1 DELTANC IN PRESENCE OF 200 MM NABR | Yes | Passing each other, Forming a complex with the RE inside |
| 6k5i | E148D/R147A/F317A MUTANT CLC-EC1 IN THE PRESENCE OF 20 MM NABR | Yes | Passing each other, Forming a complex with the RE inside |
| 6kkr | KCC1 IN KCL AND DETERGENT GDN | Yes | Parallel, Forming a complex with the RE inside |
| 6kxw | AQUAPORIN AQP7 IN BOUND TO GLYCEROL | Yes | turning with loop structure |
| 6n1g | AQUAGLYCEROPORIN AQP7 | Yes | turning with loop structure |
| 6nq0 | TPC2 CHANNEL IN THE LIGAND-BOUND OPEN STATE | Yes | Parallel, Forming a complex with the RE inside, turning with loop structure |
| 6nq1 | TPC2 CHANNEL IN THE APO STATE | Yes | Parallel, Forming a complex with the RE inside, turning with loop structure |
| 6nq2 | TPC2 CHANNEL IN THE LIGAND-BOUND CLOSED STATE | Yes | Parallel, Forming a complex with the RE inside, turning with loop structure |
| 6pis | K+ CHANNEL TRAAK (K2P4.1) - FAB COMPLEX | Yes | Parallel, Forming a complex with the RE inside, turning with loop structure |
| 6poj | AQUAPORIN 1 | Yes | turning with loop structure |
| 6qf5 | AQUAPORIN 2 CRYSTALLIZED ON A SILICON CHIP | Yes | turning with loop structure |
| 6qim | ATPIP2;4 | Yes | turning with loop structure |
| 6qzi | AQUAPORIN 7 | Yes | turning with loop structure |
| 6qzj | AQUAPORIN 7 | Yes | turning with loop structure |
| 6r7r | GLUTAMATE TRANSPORTER HOMOLOGUE GLTTK IN COMPLEX WITH D-ASPARTATE | Yes | |
| 6rv2 | POTASSIUM ION CHANNEL TASK-1 (K2P3.1) IN A CLOSED CONFORMATION | Yes | Parallel, Forming a complex with the RE inside, turning with loop structure |
| 6rv3 | POTASSIUM ION CHANNEL TASK-1 (K2P3.1) IN A CLOSED CONFORMATION WITH A BOUND INHIBITOR BAY 1000493 | Yes | Parallel, Forming a complex with the RE inside, turning with loop structure |
| 6rv4 | POTASSIUM ION CHANNEL TASK-1 (K2P3.1) IN A CLOSED CONFORMATION WITH A BOUND INHIBITOR BAY 2341237 | Yes | Parallel, Forming a complex with the RE inside, turning with loop structure |
| 6uwf | GLTPH IN COMPLEX WITH L-ASPARTATE AND SODIUM IONS IN OUTWARD-FACING STATE | Yes | |
| 6v1q | TWO-PORE CHANNEL 3 | Yes | Parallel, Forming a complex with the RE inside, turning with loop structure |
| 6v36 | K2P2.1(TREK-1)I110D APO CHANNEL | Yes | Parallel, Forming a complex with the RE inside, turning with loop structure |
| 6v37 | K2P2.1(TREK-1)I110D:RUR:ML335 BOUND CHANNEL | Yes | Parallel, Forming a complex with the RE inside, turning with loop structure |
| 6v3c | K2P2.1(TREK-1)I110D:RU360 BOUND CHANNEL | Yes | Parallel, Forming a complex with the RE inside, turning with loop structure |
| 6v3i | K2P2.1(TREK-1)I110D:RUR BOUND CHANNEL | Yes | Parallel, Forming a complex with the RE inside, turning with loop structure |
| 6wlv | TASK2 IN MSP1D1 LIPID NANODISC PH 6.5 | Yes | Parallel, Forming a complex with the RE inside, turning with loop structure |
| 6wm0 | TASK2 IN MSP1D1 LIPID NANODISC PH 8.5 | Yes | Parallel, Forming a complex with the RE inside, turning with loop structure |

(*1) The PDBTM database was searched for proteins with two reentrant loops using the keywords "0 [type] AND 2 [n_loop]", and structures were checked one by one to investigate if the reentrant loops were facing each other.
